# Integrated annotation and analysis of genomic features reveal new types of functional elements and large-scale epigenetic phenomena in the developing zebrafish

**DOI:** 10.1101/2021.08.09.454869

**Authors:** Damir Baranasic, Matthias Hörtenhuber, Piotr Balwierz, Tobias Zehnder, Abdul Kadir Mukarram, Chirag Nepal, Csilla Varnai, Yavor Hadzhiev, Ada Jimenez-Gonzalez, Nan Li, Joseph Wragg, Fabio D’Orazio, Noelia Díaz, Benjamín Hernández-Rodríguez, Zelin Chen, Marcus Stoiber, Michaël Dong, Irene Stevens, Samuel E. Ross, Anne Eagle, Ryan Martin, Pelumi Obasaju, Sepand Rastegar, Alison C. McGarvey, Wolfgang Kopp, Emily Chambers, Dennis Wang, Hyejeong R. Kim, Rafael D. Acemel, Silvia Naranjo, Maciej Lapinski, Vanessa Chong, Sinnakaruppan Mathavan, Bernard Peers, Tatjana Sauka-Spengler, Martin Vingron, Piero Carninci, Uwe Ohler, Scott Allen Lacadie, Shawn Burgess, Cecilia Winata, Freek van Eeden, Juan M. Vaquerizas, José Luis Gómez-Skarmeta, Daria Onichtchouk, Ben James Brown, Ozren Bogdanovic, Monte Westerfield, Fiona C. Wardle, Carsten O. Daub, Boris Lenhard, Ferenc Müller

## Abstract

Zebrafish, a popular model for embryonic development and for modelling human diseases, has so far lacked a systematic functional annotation programme akin to those in other animal models. To address this, we formed the international DANIO-CODE consortium and created the first central repository to store and process zebrafish developmental functional genomic data. Our Data Coordination Center (https://danio-code.zfin.org) combines a total of 1,802 sets of unpublished and reanalysed published genomics data, which we used to improve existing annotations and show its utility in experimental design. We identified over 140,000 cis-regulatory elements in development, including novel classes with distinct features dependent on their activity in time and space. We delineated the distinction between regulatory elements active during zygotic genome activation and those active during organogenesis, identifying new aspects of how they relate to each other. Finally, we matched regulatory elements and epigenomic landscapes between zebrafish and mouse and predict functional relationships between them beyond sequence similarity, extending the utility of zebrafish developmental genomics to mammals.

## Introduction

Zebrafish is used as a model vertebrate in more than 1200 laboratories worldwide (ZFIN) with dynamically increasing number of studies^1^ of organismal, cell and gene function in development, regeneration, behaviour, toxicology, and disease modelling^2^. The increasing popularity of zebrafish in biomedical research is due to its experimental advantages^3, 4^, wide ranging genetics resources (e.g. ZFIN^5^) and conservation of the genetic basis of most diseases between human and fish^6^. Most (75-80%) human genes associated with disease have at least one zebrafish orthologue, making zebrafish an attractive experimental system with convenient genetic manipulation tools^7, 8^. The number of developmental genomics studies in zebrafish is increasing, being facilitated by the third best annotated vertebrate genome. Some of the biological findings uncovered by these studies include the discovery of chromatin signatures^9–11^, DNA codes of promoter usage^12^, post-transcriptional regulation of mRNAs^13–15^, as well as the role of stem cell factors^16, 17^ and regulatory patterns of DNA methylation^18–20^ during zygotic genome activation (ZGA). In addition, zebrafish single cell genomics helped in pioneering applications for spatial resolution of developmental lineage-specific transcriptomes during development^21^. Comparative genomics applications have led to prediction of conserved regulatory elements and their long-range target genes^22^. Transgenesis can be upscaled to truly high-throughput^23^ which allowed the validation of conserved, disease associated human enhancers predicted by genomics^24, 25^. However, despite the many landmark developmental genomics studies utilizing zebrafish, this model species has lacked systematic functional annotation programmes at a scale seen in other key animal models, such as ENCODE^26^, Roadmap Epigenome^27, 28^, and modENCODE^29, 30^. While annotation of promoters and enhancers was reported for adult zebrafish muscle and brain tissues^31^, and despite the fact that embryo and larval stages represent the bulk of zebrafish-based research, these stages lack functional genomic annotation resources. The largely disparate genomics resources produced by the zebrafish community remain mostly inaccessible to the thousands of potential laboratories. Recognising this need DANIO-CODE was established as a multinational bottom-up effort of data producers^32^.

DANIO-CODE aimed for the functional annotation of the developing zebrafish genome by i) collecting all published and producing new genomics data by 38 laboratories worldwide and standardizing their unstructured metadata annotation; ii) creating and maintaining a single data coordination centre (DCC) for continued accumulation and user download of zebrafish genomics datasets^33^; iii) developing standardized analysis pipelines and remapping all sequencing datasets according to ENCODE^34^ and FANTOM Consortium standards^35^; and iv) generating an integrated track hub that allows visualization with common genome browser tools such as UCSC Genome Browser^36^, GBrowse^37^ and Ensembl^38^. Additionally, DANIO-CODE aimed to conduct the first integrated analysis of these datasets to promote discovery, functional element classification and determination of features of developmental dynamics. Finally, in this study we present novel approaches for comparative analysis of zebrafish and mammalian genomic datasets to understand the degree of synteny and conservation of genomic architecture, epigenetic and chromatin configuration, and topology among vertebrates and to expand the utility of zebrafish developmental genomics resources.

## Results

### Comprehensive collection and annotation of zebrafish developmental genomics data in the DANIO-CODE Data Coordination Center

We have established a Data Coordination Centre (DCC) protocol^33^, which was adopted and populated by the DANIO-CODE international network (https://www.birmingham.ac.uk/generic/danio-code/partners/index.aspx) by deposition of zebrafish developmental genomics data including standardised annotation of metadata of diverse, often inconsistently annotated published datasets (www.danio-code.zfin.org, **Figure 1a**). The DCC is accessible from the Zebrafish Information Network (ZFIN) and includes details about the datasets and their underlying samples and sequencing protocols using ZFIN and ENCODE nomenclature. It provides a platform for global data reprocessing for future data producers. To identify and analyse the developmental dynamics of genomic features, direct comparison across datasets produced by different laboratories and different protocols is necessary. To this end, we carried out consistent and standardised reprocessing starting from the raw sequencing data (**Figure 1a**). Raw sequencing data were collected and reprocessed for integrated analysis by standardised pipelines of ENCODE for ChIP-seq and ATAC-seq^34^, FANTOM for CAGE-seq^35^ and for Hi-C and 4C-seq, data producer pipelines were used (for details see Methods). Novel and revised pipelines are available on GitLab (https://gitlab.com/danio-code). The DCC data include 1,438 published datasets, which were contributed by data producers directly or were collected by data annotators from the DANIO-CODE consortium to complement the user-supplied data with datasets strategically selected for developmental data from the public domain. In addition, 366 datasets were generated for this study by consortium members to fill data gaps and to aid functional annotation and functional element characterisation, including 15 CAGE-seq, 18 ChIP-seq, 11 ATAC-seq, 2 HiC, and 320 4C-seq datasets (**Figure 1b**, **Suppl. Figure 1a**). The DCC thus contains 1,804 zebrafish developmental genomics datasets with 71 series across 38 developmental stages (**Suppl. Figure 1d**) and 34 tissues from 20 assay types. Breakdown of the datasets according to data types and stages of development is presented in **Figure 1b**. ChIP-seq data are presented as merged sets including 4 histone modification marks and 39 datasets for Pol II, CTCF, and transcription factors (**Suppl. Figure 1b**). The status of published and novel data collection from 38 laboratories worldwide is indicated in **Suppl. Figure 1c** and **Suppl. Table 1**. Quality checks and data comparability analyses were carried out for datasets within a data type obtained from multiple laboratories, particularly affecting RNA-seq (**Suppl. Figure 2b**), ChIP-seq (**Suppl. Figure 2d-f**), CAGE-seq (**Suppl. Figure 4**), and ATAC-seq (**Suppl. Figure 2c**) data. The DCC continues to be periodically updated (**Suppl. Figure 1d**), and is openly accessible to the user community for downloading data and for uploading new datasets.

**Figure 1.**
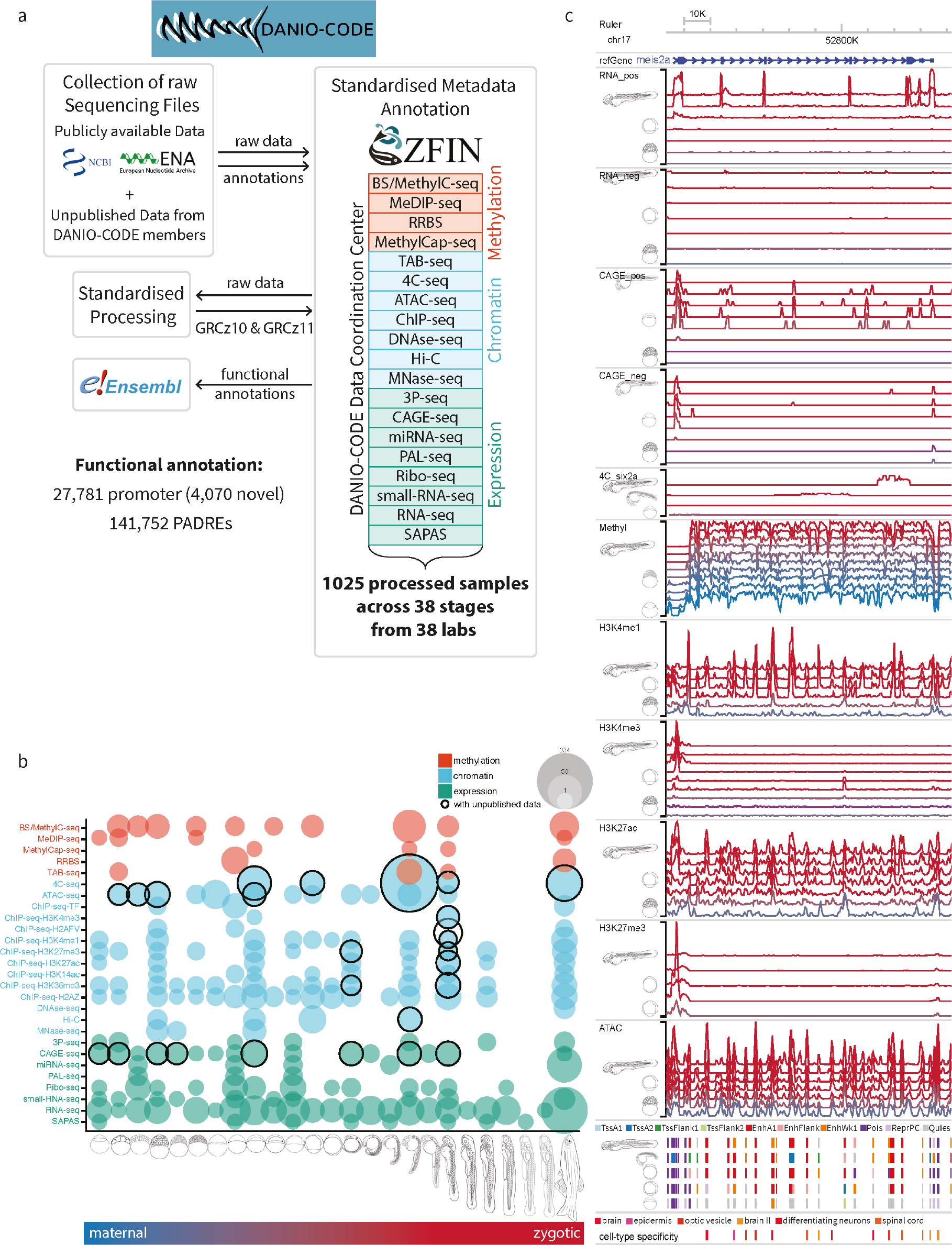
Comprehensive collection and annotation of zebrafish developmental genomics data. **a**, Collection and manual annotation processes of datasets with input and output interactions with the DANIO-CODE Data Coordination Center. **b**, Components of the open repository for developmental multi-omics data for zebrafish with data type on the Y axis and developmental stage represented on the X axis of the matrix. Unpublished data first reported in this study are highlighted with black circles. **c**, Visualisation of temporal dynamics of selected epigenomic features during development at a developmentally active locus. Colouring of tracks represents developmental series from maternal (blue) to zygotically active stages of embryogenesis (red). Symbols indicate representative stages.

The resulting data and reprocessed multi-omics datasets represent the first comprehensive annotation of the zebrafish genome during normal embryonic development. It is available as an annotated track hub among the public track hubs in the UCSC browser (genome.ucsc.edu) and uploadable in Ensembl genome browser. **Figure 1c** provides an example track set at a developmentally-regulated locus covering selected stages from fertilisation to the Long-pec stage visualised by the Washington University Epigenome browser^39^, where tracks have been modified to allow simultaneous visualisation of the developmental time. The bulk data-based tracks include not only gene expression and epigenomic features, but also new annotation of approximately 140,000 Predicted ATAC-seq-supported Developmental Regulatory Elements (PADRE) annotated by ChromHMM categories and assigned to open chromatin regions (**Figure 1c**). The bulk data-based predictions for regulatory elements are complemented with cell-type specificity of candidate regulatory elements generated by intersecting them with single cell ATAC-seq-based candidate regulatory element annotations^40^.

### Annotation of transcribed elements and characterisation of core promoters

As genome-wide transcriptome analyses^41–44^ fail to annotate 5’ UTRs precisely, we used DANIO-CODE expression data to correct current Ensembl versions of developmentally-active gene and transcript models. We utilised 139 developmental RNA-seq samples to identify 30,832 genes comprising 51,033 transcripts (**Figure 2a**, **Supplementary Table 2**), among them 131 novel transcripts of uncertain coding potential (TUCP) and 251 lncRNA genes that were not previously annotated by Ensembl, but supported by CAGE signals (**Suppl. Figure 3**). To detect the 5’ end of transcripts precisely, we mapped transcription start sites (TSSs) from 34 CAGE samples in 16 developmental stages (**Figure 2a**). We applied standardised filtering and promoter-calling criteria to CAGE datasets produced by distinct technologies (Methods, **Suppl. Figure 4a-c**), which on average yielded 22,500 active promoters per CAGE sample (**Suppl. Figure 4d**, **Suppl. Table 4**), thus adding 4,070 novel promoters located at least 1kb apart from an Ensembl transcript start to 18,646 previously annotated Ensembl TSSs (GRCz10). Together, the called promoters correspond to 16,303 genes (**Figure 2a**) with 3,930 Ensembl transcripts being associated with more than one alternative promoter region.

**Figure 2.**
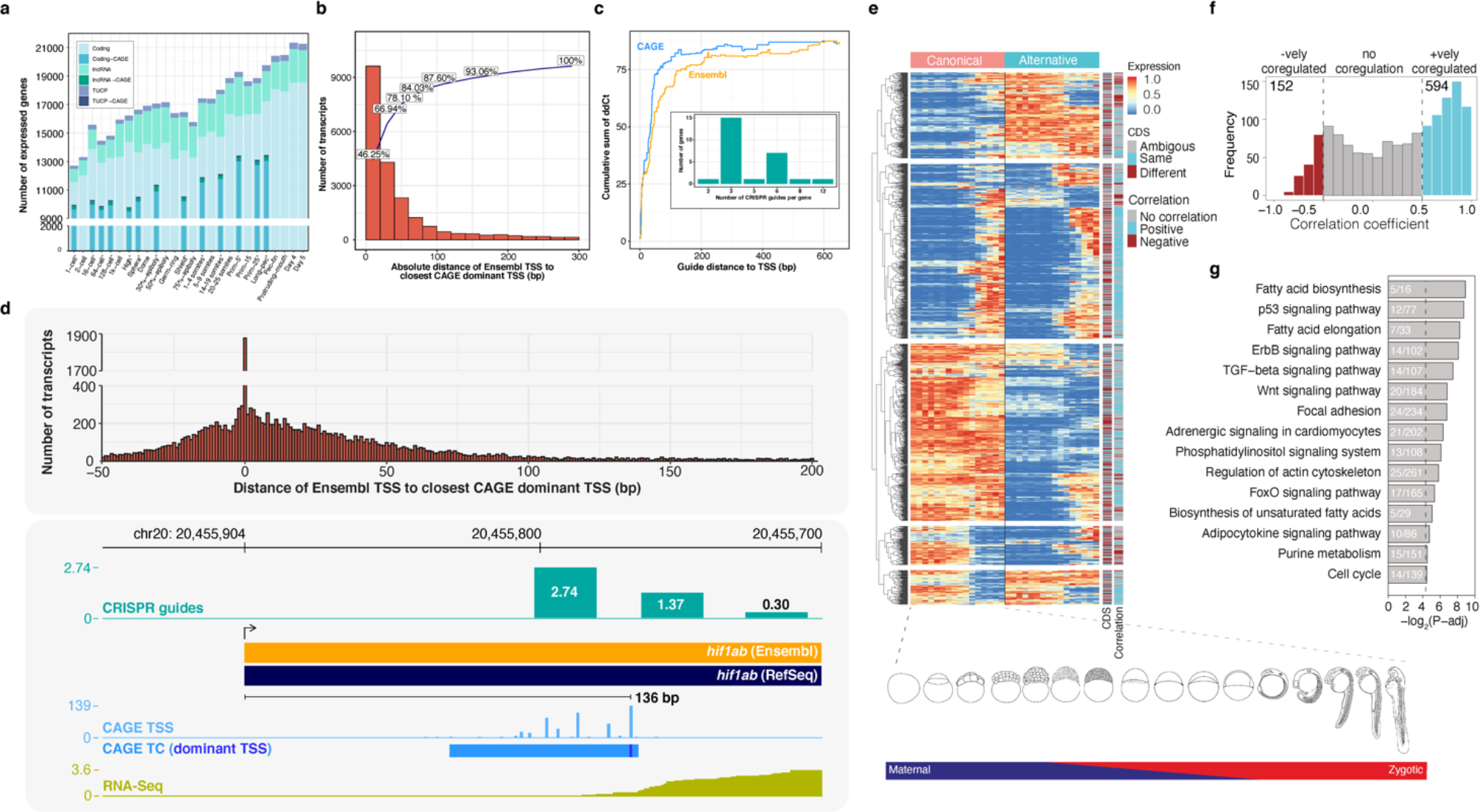
Transcript categories and single nucleotide resolution 5’ end verification during development. **a**, DANIO-CODE transcript 5’ ends supported by CAGE TSS during stages of development. **b,** Distribution of absolute distance of Ensembl TSSs to CAGE dominant TSSs in the Prim-5 stage. **c**, Relationship between guide distance to TSSs and ddCt, highlighting the CAGE precision over Ensembl. Insert: number of dCas guides for all 26 tested genes. **d,** CAGE defined TSSs increase the accuracy of promoter identification and support dCas inhibition guide reagent designs. **e**, Heatmap shows the dynamics of expression levels of canonical and alternative promoters across 16 developmental stages represented as images. Expression levels are scaled in the range of 0-1 for each row. Canonical and alternative transcripts using same and different CDS start are denoted as same and different respectively. Either of the transcript pairs that are non-coding or without full CDS annotation are denoted as ambiguous. **f,** Distribution of correlation coefficient of expression levels of canonical and alternative promoters across 16 developmental stages. **g,** Enrichment of KEGG pathways on multi promoter genes. The adjusted p-value cut-off is 0.05, denoted by a vertical dashed line. The number of genes in KEGG pathways and those overlapping with multi promoter genes are shown inside bars.

Our above definition of single nucleotide-resolution promoter may offer important guidance for promoter-targeted gene manipulation to block transcription. Transcription block is necessary to avoid genetic compensation by transcriptional adaptation, which may mask the phenotypes in reverse genetic experiments that inactivate the coding sequence^45^. As such inhibition of transcription requires precise annotation of promoters, we compared Ensembl’s RNA-seq-based TSS with our CAGE-seq-based TSS and found a substantial distance discrepancy in thousands of genes (**Figure 2b**). In the majority of promoters (>55%), the CAGE-defined main TSS falls >20 bp outside of the 5’ end of the Ensembl (GRCz10) gene model, and for >20% of genes the difference is >40 bp at the Prim-5 stage, potentially impacting on guide-RNA design for CRISPR/Cas targeting (**Suppl. Fig. 5a**). To test the impact of guide design position in relation to the dominant TSS, multiple guide positions were designed and tested for 26 genes. Impact of dead Cas system (dCas) inhibition on transcript levels decreased with increased distance between the guide target and the dominant CAGE-defined TSS. Efficiency of dCas inhibition was higher when CAGE-dominant start sites were used as compared to Ensembl start sites (**Figure 2c, d, Suppl. Table 5**) demonstrating the importance of TSS detection accuracy in transcription inhibition and the improved accuracy of CAGE over Ensembl in promoter detection.

We systematically analysed the 5’ end of Ensembl and RNA-seq transcripts supported by CAGE and identified 1,293 multi-promoter genes (**Suppl. Table 6**). Out of these, 1,176 multi-promoter genes had assigned one canonical promoter and one alternative promoter (see Methods), while 117 multi promoter genes had one canonical promoter and two or more alternative promoters. The large majority of these genes (929) have human orthologs that also have alternative promoters. Correlation of expression levels of canonical and alternative promoter pairs indicate both convergent (cyan in **Figure 2e, f**) and divergent (brown) dynamics during embryonic development. The expression of canonical promoters is on average higher than those of alternative promoters (**Suppl. Figure 5b**). Among 978 transcript pairs with full-length coding sequence (CDS) annotation, 373 (38%) of the alternative promoters affected only the 5’ UTR (e.g., *dag1*, **Suppl. Figure 5c**), whereas the remaining 605 reside downstream of the start codon and alter the N-terminus of the protein sequence (e.g., *bmp6*, **Suppl. Figure 5d**), highlighting the functional importance of alternative promoter annotations. Notably, the 5’ UTRs of alternative transcripts are significantly shorter when they utilize the same start codon as the canonical transcript, whereas there was no significant difference in 5’ UTR length when they utilize different start codons (**Suppl. Fig. 5f**) suggesting distinct 5’ UTR regulation between alternative promoters. Compared to an average of approximately 4 promoters per gene across human tissues and cell types^35^, alternative promoters were less frequently observed during zebrafish embryonic development. However, the multi promoter genes in zebrafish are enriched in multiple KEGG signalling pathways (e.g., p53, ErbB, TGF-beta and Wnt signalling, **Figure 2g**), many of which are dysregulated in cancers^46^, suggesting deep evolutionary conservation of alternative promoters in signal transduction-associated genes in vertebrates.

### Classification of the genomic regulatory regions in development

Next, we aimed to generate the first comprehensive atlas of developmental regulatory elements for zebrafish in which we classify and characterise their developmental dynamics. We defined *cis*-regulatory elements (CREs) as ATAC-seq^47^ peaks reproducible between replicates, and termed them Predicted ATAC-seq-supported Developmental Regulatory Elements (PADREs), that we further classified based on presence of CRE-associated histone modification marks (H3K4me3, H3K4me1, H3K27ac, and H3K27me3) with ChromHMM^48, 49^ (**Figure 3a**, **Suppl. Fig. 6a-c**). We defined PADREs in 4 pre-ZGA and 7 post-ZGA stages (**Suppl Figure 7a**), and classified them with ChromHMM in 5 post-ZGA stages.

**Figure 3.**
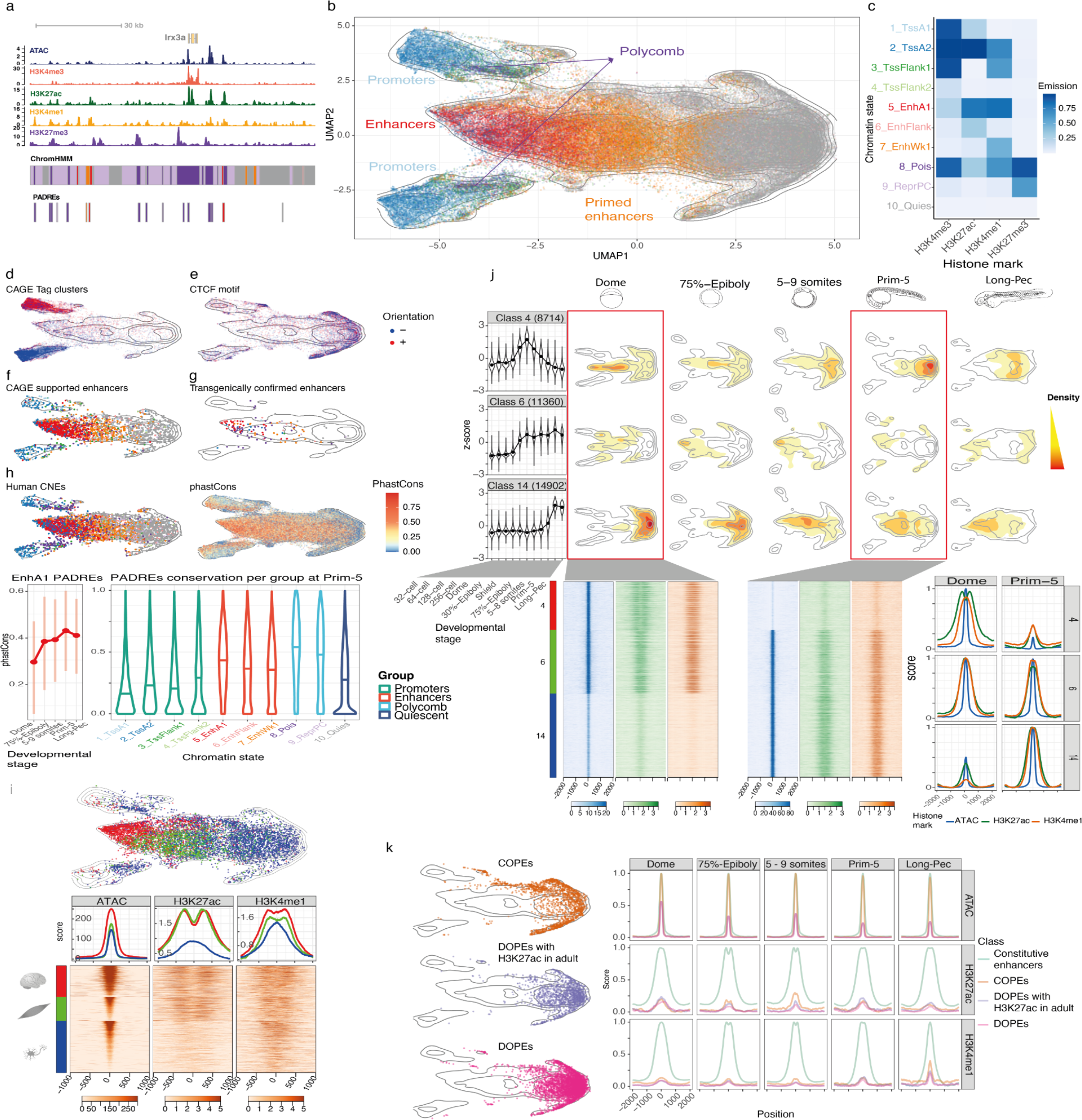
Classification of developmental cis regulatory elements. **a**, Genome browser screenshot showing ChromHMM classification of PADREs, and respective histone post-translational modification signals used to define them. **b,** UMAP plot of PADREs at the Prim-5 stage. Each point represents one open chromatin region, coloured by functional assignment. **c.** Occurrence probabilities of chromatin marks for ChromHMM states. The states function was manually assigned using The Roadmap Epigenomic annotations as reference. 1_TssA1, 2_TssA2 = Active TSS, 3_TssFlank1, 4_TssFlank2 = TSS Flanking region, 5_EnhA1 = Active enhancer, 6_EnhFlank = Weak enhancer, 7_EnhWk1 = Primed enhancer, 8_Pois = Poised elements, 9_PcRep = Polycomb repressed regions, 10_Quies = Quiescent state. **d-g.** UMAP plot showing PADREs overlapping with CAGE promoters (**d**), CTCF motif (**e**), eRNA enhancers (**f**), and transgenically validated enhancers (**g**). **h,** UMAP plot showing the mean phastCons score for each PADRE (top right) and overlap with human CNEs (top left). The bottom sub-panel shows the distribution of tCons scores of active enhancers throughout development (left), as well as the distribution of the phastCons score for PADREs separated by function at the Prim-5 stage (right) **i,** Top: Position of cell type-specific elements on the UMAP plot. Bottom: ATAC, H3K27ac, and H3K4me1 signals around the peak summit of cell-type specific PADREs. **j,** Upper: Openness profile of selected SOM classes (4: early, 6: post-ZGA constitutive, and 14: late class), and their position density on the UMAP plots of different developmental stages. Bottom: Heatmap of signal intensity of ATAC, H3K27ac, and H3K4me1 at the Dome and the Prim-5 stages, along with their respective profiles. **k,** left: Position of COPEs, DOPEs, and DOPEs marked with H3K27ac in adult tissues on the UMAP plot. Right: Profiles of ATAC, H3K27ac, and H3K4me1 of COPEs, DOPEs, DOPEs marked in adult tissues, and other constitutive elements throughout development.

To visualise and monitor the dynamics of PADREs across early development, we developed a UMAP-based method (Methods, **Suppl. Figure 8a, b**). We hypothesized that UMAP dimensionality reduction of the epigenetic landscape at open chromatin regions will predict their classes and potentially highlight novel functional subclasses during development. The UMAP plot for Prim-5 PADREs (**Figure 3b**) demonstrates separation of most ChromHMM functional classification groups including separation of promoters from enhancer classes (**Figure 3c** and **Suppl. Fig 7d**). Near-symmetry on the y-axis reflects DNA strand directionality and is most prominent among promoters. The UMAP classification of promoters by position and orientation was verified by plotting CAGE-seq defined promoters on the UMAP (**Figure 3d**). To explore UMAP clusters further, we focussed on PADREs that do not overlap with any chromatin marks. They form a large bulk of clusters on the right, flanking two prominent clusters, which stretch upward and downward from the right apex. These domains of the UMAP are enriched for the binding motif of CTCF with well-positioned flanking nucleosomes^50^ (**Figure 3e**, **Suppl. Figure 9**). Enhancer predictions were functionally validated with two independent sets: 1) by plotting enhancers characterised with bidirectional eRNA signals^51^ called from newly generated nuclear CAGE data; and 2) by plotting a manually curated catalogue of published enhancer elements, which were identified and functionally validated in transgenic reporter assays (**Suppl. Table 7**). Both sets overlap highly with enhancer-classified PADREs and co-localise with active enhancers on the UMAP plot (**Figure 3f, g Suppl. Figure 7c, d**), indicating the utility of the UMAP method for enhancer detection. DNA Methylation analysis of PADREs revealed that PADREs are either persistently hypomethylated across stages or are gradually hypermethylated during development before becoming hypomethylated in adult somatic tissue. PADREs with persistent hypomethylation are CG-rich and are associated with regions such as the TSS, while the dynamically methylated PADREs are less CG-dense and associated with elements such as enhancers. Additionally, the different classes of dynamically methylated PADREs vary in the onset and degree of their hypermethylation and hypomethylation events. For example, the enhancer class A is the first to begin hypomethylating at 24hpf and are made up of previously identified conserved phylotypic enhancers^18^ (**Suppl Fig 7f**).

Next, we assessed the evolutionary conservation of PADREs by overlapping with human Conserved Non-Coding Elements (CNEs) and calculating the phastCons score for each PADRE (**Figure 3h**, top). Early acting enhancers appear less conserved than the later functioning ones in both analyses (**Figure 3h**, bottom left; **Suppl. Figure 7e**). phastCons scores of enhancers are higher on average than promoters (P-value < 2.2e-16) (**Figure 3h**, bottom right). Poised elements are the most conserved (P-value < 2.2e-16), strongly suggesting that Polycomb-bound enhancers are a specific class of enhancers under high purifying selection, playing a critical role in differentiation and organogenesis^24, 52^, and contributing to the hourglass model of development^53^.

To assign cell type-specificity to PADREs we exploited single cell ATAC-seq^40^ data at Prim-5. UMAP plotting (**Figure 3i**, right) revealed remarkable differences in signal profiles between brain and muscle elements, which carry comparable levels of H3K27ac and H3K4me1, but distinct profiles of openness (**Figure 3i**, bottom). In contrast, differentiating neurons show a 3-fold decrease of the H3K27ac signal (**Figure 3i**, left).

Next, we sought to understand the temporal dynamics of PADREs. All stage-specific PADREs were merged to form consensus PADREs (cPADREs), containing ∼140k regions open in at least two neighbouring stages (**Suppl. Figure 7a**). We then clustered non-promoter cPADREs based on their chromatin accessibility into self-organising maps (SOMs) (**Suppl. Figure 10**). UMAP locations of 3 out of 16 SOM clusters demonstrate developmental chromatin changes associated with PADRE dynamics (**Figure 3j**, top). These clusters feature elements active early and subsequently decommissioned (Class 4), active constitutively from ZGA onwards (Class 6), and late elements (Class 14), which transition from primed to active state. We next explored the underlying changes in chromatin profiles by plotting signal heatmaps around ATAC-seq peaks (**Figure 3j**, bottom). Concordant to the distinct UMAP plots, early and late elements show different epigenetic profiles in their respective active states. Unlike late and constitutive elements, early elements were depleted of the H3K27ac mark at the peak. These findings indicate that early and late enhancers represent two different chromatin states with distinct conservation profiles.

Finally, we explored the dynamics of a set of elements we called “quiescent” state PADREs, which we identified as unmarked at any stage of development. 2,109 such regions are constitutively open throughout development without observable CRE-associated chromatin marks (**Suppl. Figure 11**). We collectively termed these elements Constitutive Orphan Predicted Elements (COPEs). They colocalise with constitutive SOM class 6 and 40% of them contain CTCF sites (**Figure 3k**, top; compare to **Figure 3e**, **Suppl. Figure 10**). In contrast, another non-marked open chromatin set (11,044) is open only in specific developmental stages and called Dynamic Orphan Predicted Elements (DOPEs, **Figure 3k**, **Suppl. Figure 11a**). Using data from^31^, we found 2513 DOPEs that contain active chromatin marks in adult tissues, revealing the existence of novel types of regulatory elements whose chromatin opening precedes enhancer-associated histone mark deposition.

### Chromatin architecture classification and developmental specialisation of Pol II gene promoters

To reveal any developmental regulatory principles, we exploited the chromatin features of PADREs to functionally classify Pol II promoters. First, we characterised these promoters from the Dome and Prim-5 stages based on their chromatin accessibility at nucleosome-resolution, revealing 8 clusters (**Figure 4a, Suppl. Fig. 12, Suppl. Table 8**). The clusters differ mostly in their upstream configuration, including the width of the nucleosome-free region (NFR), the signal strength of the NFR, and the presence of optional upstream open regions (**Figure 4a**), which follows GC content (**Suppl. Figure 12a**). These NFR variations are characterised by mark presence and distinct pattern of upstream opposite strand transcription (e.g., *upstream offset* and *double NFR* promoters) with distinct distances between the main TSS and flanking nucleosomes (e.g., wide and strong open or *narrow* and *medium constitutive*) and TSS profiles (**Suppl. Figure 12b**). These classes also show notable differences in H3K4me1, H3K4me3 and H3K27ac histone modification patterns (**Figure 4b**), confirmed also by the differing UMAP positions of promoter PADREs (**Figure 4c**). Apart from *weak open,* each class produces antisense transcription (prompts REF), including *Double NFR*, *wide* and *upstream offset* classes, which show CAGE expression from both the main NFR and another upstream region, with the sense transcription being stronger than antisense (**Figure 4a**). Notably, the architecture classes remain stable over developmental time (**Figure 4d, Suppl. Figure 12c**), suggesting they represent distinct regulation mechanisms acting on the genes rather than reflecting stage-dependent gene activity states. *Wid*e and *strong open* classes contain the most conserved promoters (**Figure 4e, Suppl. Figure 12d**), and are enriched in transcription regulator genes (**Figure 4g, Suppl. Figure 12e**). However, the promoter classes show distinct developmentally dynamic temporal expression (**Figure 4f, Suppl. Figure 12f**) with notable enrichment of the *double NFR* class for maternally expressed genes in contrast to the predominantly early and late zygotic *weak open* and *medium zygotic* classes, respectively. The promoter classes also show distinct GO enrichment categories (**Figure 4g**). Overall, our approach offers the first promoter architecture classification for zebrafish and indicates functional specialisation of promoter classes.

**Figure 4.**
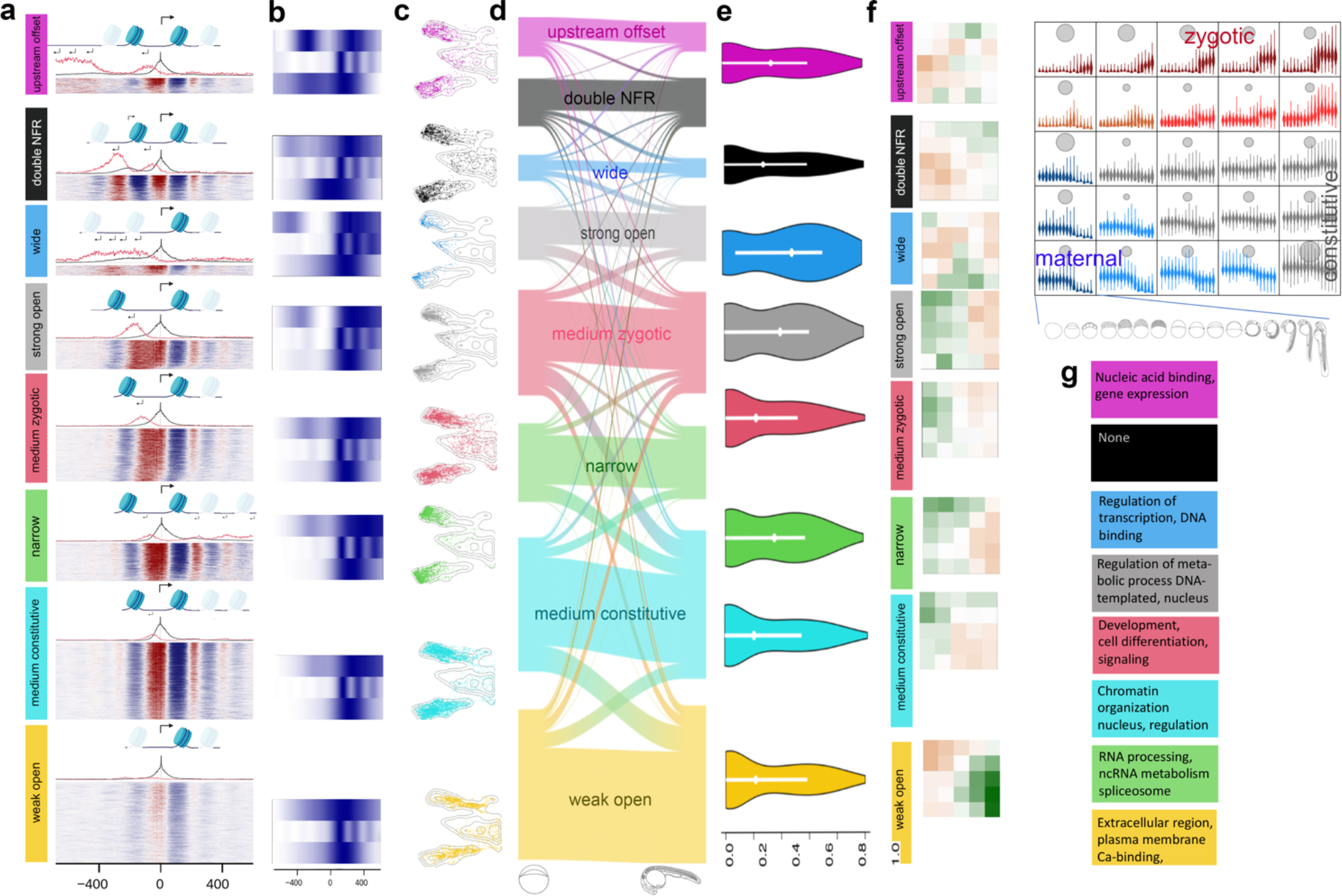
Chromatin architecture classification and developmental specialisation of Pol II gene promoters. **a**, Heatmap of chromatin accessibility profiles aligned to dominant TSS per promoter at the Prim-5 stage. Nucleosome-free regions (red) are superimposed with nucleosome positioning (blue). Stack height reflects number of promoters. Above each heatmap, combined histograms of CAGE expression are shown. Black, forward TSSs, red, reverse orientation TSSs (The scale is amplified 10x in relation to forward transcription). Nucleosome positioning is symbolised above alignments and black arrows indicate transcription direction; size indicates relative strength. Promoter configuration classes are colour-coded consistently in all panels including **Suppl. Figure 12**. **b**, Aggregated H3K4me1, H3K4me3, H3K27ac (top to bottom) ChIP-seq signals for classes as in **a** are aligned to dominant TSS. **c**, UMAP profiles of promoter classes at the Prim-5 stage. UMAPs are cropped to highlight promoter PADREs. **d**, Flow diagram indicates the relationship between promoter configuration class at the Dome stage (left edge, **Suppl. Figure 12**) and the Prim-5 stage (right edge). Band width represents the number of promoters. **e**, Violin plot of phastCons vertebrate conservation distribution of promoters. Each class is aligned to **a**. **f**, Classification of promoter expression during development with self-organising maps (SOM). On the top right, 5×5 diagrams contain violin plots with stage-by-stage expression levels. Blue to red spectrum indicates maternal to zygotic expression dynamics of promoter clusters. Surface areas of grey circles indicate the number of promoters per cluster. Stages of development are symbolised below the SOM array. On the left, the mustard - positive green-negative colour spectrum in SOMs indicates the enrichment in promoter overlap between promoter expression classes (SOM) in each chromatin architecture class **a**. **g,** Enriched GO categories for each promoter architecture class. Full GO lists are in **Suppl. Table 9**.

### Temporal dynamics and locus organisation of enhancers during development

Key genes regulating development are under the control of numerous long-range enhancers, which often overlap with highly Conserved Noncoding Elements (CNEs). Main features of those regulatory landscapes are captured by the genomic regulatory block (GRB) model^22^, in which a target gene integrates the input of multiple enhancers spanning hundreds of kilobases. The region typically contains other “bystander” genes that do not respond to those enhancers. The extent of GRBs coincides with those of topologically associating domains (TADs) around developmental genes^54^ (**Figure 5a**). We exploited DANIO-CODE functional annotations to characterise chromatin opening and interaction topology in those extremely conserved, yet poorly understood loci and to understand their regulatory role in gene expression in their respective TADs.

**Figure 5.**
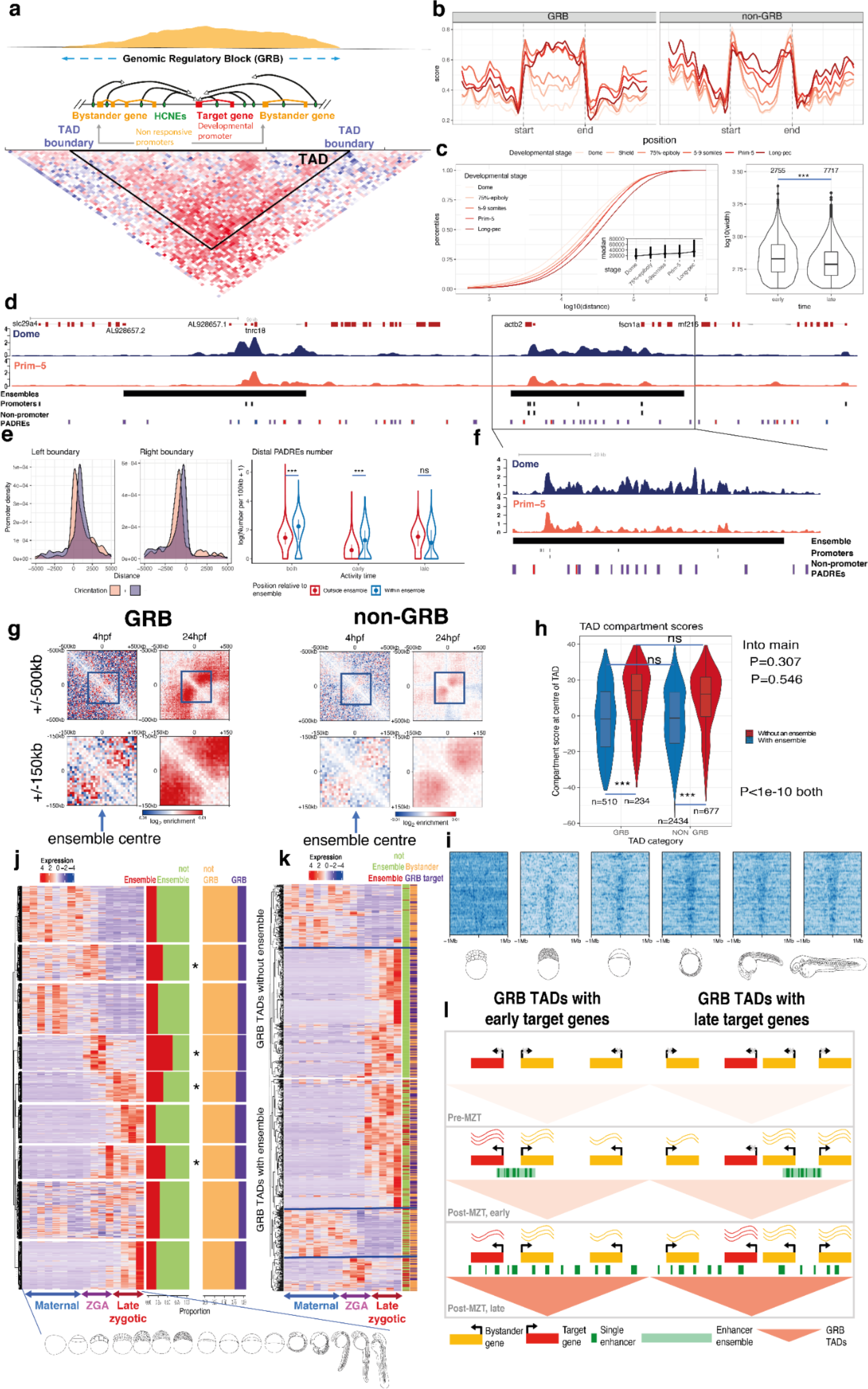
Dynamics and function of open chromatin and H3K27ac topology organisation on early embryo development. **a**, Schematic representation of genomic regulatory blocks (GRBs). Middle: basic components of a GRB. GRB Enhancers (green) regulating the target genes span the entire length of the GRB. Top: typical density pattern of conserved non-coding elements in a GRB, most of which overlap enhancers. Bottom: HiC contact matrix within a GRB. **b,** Chromatin opening profiles through developmental stages in TADs (length normalised). Score values were normalised to a 0 - 1 scale for comparison. **c** Left: Distance distribution of enhancer-associated PADREs to the closest promoter within GRB TADs. Right: Width distribution of concatenated enhancer-associated ChromHMM segments. Singletons shorter than two bins (400 bp) were excluded. The number of segments is shown above each violin plot. **d,** Genome browser view of a GRB TAD showing H3K27ac signals in the Dome and Prim-5 stages, H3K27ac ensembles (annotated by black bar), CAGE promoters (black blocs), and non-promoter PADREs. Blue PADREs represent one active in the Dome stage, the red PADREs are active in the Prim-5 stage, and purple PADREs are active in both stages. **e**, left, Density of CAGE promoters on ensemble boundaries. Right, the number of non-promoter PADREs per 100kb in TADs containing ensembles. The x-axis shows the developmental stage in which the PADRE is H3K27ac marked (early, late, or both). The location of promoters in respect to the ensemble is shown in different colours. **f,** left: A zoomed in genome browser view of a H3K27ac ensemble. The labels are the same as in d. **g,** Aggregate contact enrichment centred on ensembles at stages as indicated. Blue squares are magnified at the bottom. **h,** TAD compartment score distribution. **i,** Heatmaps of H3K27ac signal across GRB TADs containing ensembles through developmental stages. TADs are ordered by their width in descending order and fixed on the TAD centre. **j,** CAGE expression patterns of gene classes separated by SOM. Bar plots in the middle show the proportion of ensemble-associated and GRB genes in each class respectively. **k,** Gene expression pattern of GRB target and bystander genes. The left sidebar shows an ensemble association for each gene. The right sidebar shows the target or bystander assignment for each gene. **l,** A model describing the influence of an H3K27ac ensemble on expression of GRB target genes. If the H3K27ac ensemble is in contact with the target gene, it can be expressed early on. If the target gene is not in contact with the H3K27ac ensemble, it can only become expressed once the long-range interactions are present.

For our analyses, we distinguish *GRB TADs,* characterised by a high density of extreme non-coding conservation, from *non-GRB TADs* (see Methods). In the regions corresponding to late (Long-pec) embryo TADs, the chromatin starts opening at the boundaries, as early as in the Dome stage, and remains open throughout the analysed stages (**Figure 5b, Suppl. Figure 13a**). GRB-associated TADs show a strong increase in accessibility across the entire TAD, whereas in non-GRB TADs the increase in accessibility is mild and occurs later (**Figure 5b**). Thus, establishment of the GRB TADs might guide the distribution of long-range enhancers within them. TADs start to form early on but form fully only at later developmental stages^55, 56^ (**Suppl. Figure 13c**). By comparing early- and late-stage TADs, we showed that promoter-regulatory elements interactions are generally shorter in early-stage TADs (18kb median distance in the Dome stage vs 34kb median distance in the Long-pec stage; **Figure 5c**, left), which suggests that early expressed genes rely less on long-range interactions within TADs.

When we estimated the potential activity of enhancer candidates by H3K27ac occupancy in TADs, we observed that PADREs marked by H3K27ac or H3K4me1 in late developmental stages are numerous, short, and distributed throughout the entire TAD length. Contrary to that, PADREs active in the Dome stage are rare and often occur in large continuous, uninterrupted H3K27ac clusters at the Dome stage (**Figure 5c**, right; **Suppl. Figure 13b, Figure 5d**). We detected ∼1600 such clusters with the ROSE algorithm^57^, of which ∼1300 fall in TADs and are specifically enriched in GRB TADs (**Suppl. Figure 13d**). These clusters are reminiscent of super-enhancers^57, 58^, although they are more frequent than 231 reported in mouse^57^ and 411 in zebrafish^56^. Given the unusual scale and distinct early appearance before lineage determination (when previously reported super-enhancers appear), we distinguish these regions from super-enhancers and call them *H3K27ac ensembles*. We hypothesized that the formation of these ensembles might be associated with lack of strong TADs in early stages, and may reflect active regulatory elements clustering at a shorter range around early active promoters. To test this, we investigated the relationship between distribution, interactions between PADREs and H3K27ac ensembles and associated gene activity during early and late embryogenesis.

We found that promoters appear to be enriched at the boundaries of the H3K27ac ensembles (**Figure 5e**, right) and that the ensembles contain a majority of candidate enhancer PADREs detected in early stages (**Figure 5e**, left). In contrast, PADREs that are active only later in development and likely represent long range regulators, which appear along the entire TAD (**Suppl. Figure 13b**), are not selectively enriched in ensembles (**Figure 5e**, left). PADREs residing in H3K27ac ensembles active only early are less evolutionarily conserved than either the early ones that remain open or the ones that appear later, in agreement with the hourglass model of conservation of embryonic development^53^, and highly conserved elements are neither depleted, nor enriched in ensembles (**Suppl. Fig 13e**). Moreover, the H3K27ac mark present along the entire length of the ensemble becomes restricted to individual H3K27ac peaks associated with PADREs by the Prim-5 stage (**Figure 5f**).

Consistent with our hypothesis of the potential H3K27ac ensemble role in early gene regulation, we observed increased contacts within them at the Dome stage by Hi-C analysis in both GRB and non-GRB TADs. At the Prim-5 stage, strong contacts spread throughout the entire TAD with H3K27ac e (**Figure 5g, Suppl. Figure 14**). TADs with H3K27ac ensembles present at Dome stage generally belong to the A-compartment at the Prim-5 stage (**Figure 5h**), arguing for a role for H3K27ac ensembles in timely opening of chromatin in their host TADs. Indeed, in GRB TADs, the H3K27ac mark propagates from H3K27ac ensembles to fill the entire TAD in later stages (**Figure 5i**).

To examine how H3K27ac ensembles may influence gene expression, we classified all promoters within TADs by their expression dynamics using SOM into 9 clusters (**Figure 5j** left). H3K27ac ensemble-associated promoters mostly sequester into clusters with highest expression in early post-ZGA stages (**Figure 5j** right with asterisk). However, promoters within GRBs (20% of analysed promoters) are enriched in clusters of late-expressed genes, as expected for key lineage-determining and other developmental regulator genes associated with GRBs. In GRB TADs without an early ensemble, two classes of genes are present: ubiquitously expressed (GRB bystanders), and late zygotic expressed (likely GRB target genes). In GRB TADs with an ensemble, we observe clusters of genes peaking in early post-ZGA stages, suggesting that ensembles assist in activation early acting developmental genes, but not the late acting GRB target genes (**Figure 5k**, modelled in **Figure 5l**).

### Similarities of epigenomic features between fish and mammals suggest functional conservation of intraTAD epigenetic subdomains and associated enhancer topology organisation

Next, we investigated whether the functional annotation of non-coding elements in the developing zebrafish could be exploited for the discovery of conserved epigenomic features shared among vertebrates, which may aid in predicting functional conservation of CREs in shared epigenetic domains. Existing comparative epigenomic methods rely on direct alignments between species of interest^59, 60^. However, the large evolutionary distance between fish and mammals limits the power of comparison due to loss of sequence similarity of potentially functionally equivalent elements. To address this, we developed a method to predict functional conservation across large evolutionary distances and genomic scales without direct sequence alignments, making use of the fact that functional elements often maintain collinear syntenic positions, while they scale with genome size between species, particularly in TADs with GRBs^22, 54, 61, 62^. We exploited the numerous high-quality reference genomes and used them as bridging species to identify regions of low-level sequence conservation that may escape detection by direct sequence alignments. We established multiple reference points (multi-species anchors; **Figure 6a**) between genomes using a stepped pairwise sequence alignment approach in bridging species (**Suppl. Figure 15** and Methods), which allowed us to transform coordinates between genomes of varying sizes. Our approach also incorporates a synteny constraint, which helps minimize incorrect alignments to non-syntenic regions.

**Figure 6.**
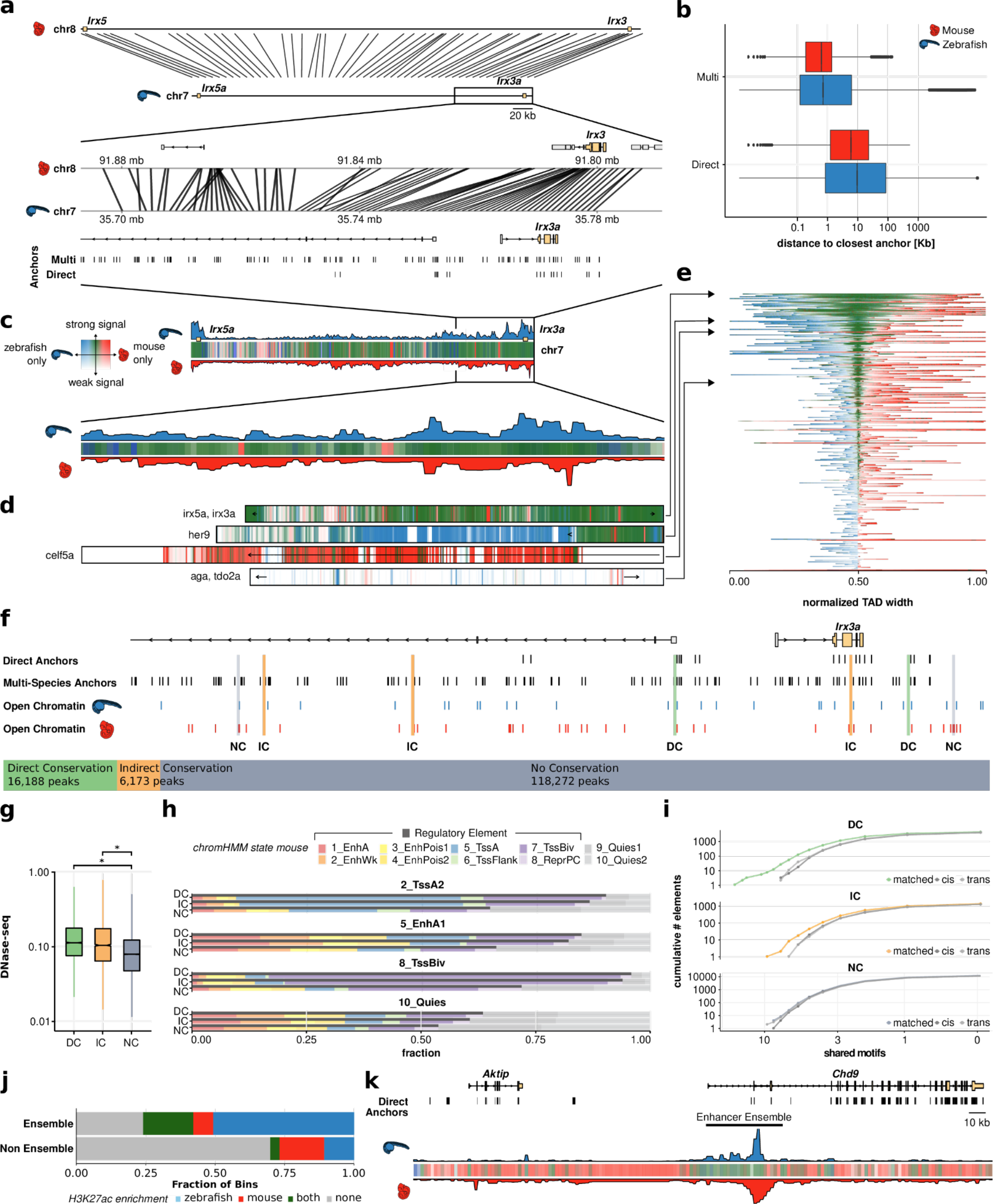
**a**, Cross-species comparison of the TAD comprising the genes Irx3/5(a) between zebrafish and mouse and a zoom-in on the locus around Irx3(a). Connecting lines represent projections of bin centers from zebrafish to mouse. **b,** Distribution of distances from the bin centers to their closest anchors in zebrafish (blue), and from their projections to their closest anchors in mouse (red), using the direct and the multi-species projection approach. **c,** Epigenetic comparison of the Irx3/5(a) TAD. H3K27me3 overlap in mapped regions is indicated as colored bars (green - mutually enriched, blue - zebrafish specific enrichment, red - mouse specific enrichment, see Methods). Opacity reflects signal amplitude and is proportional to the maximum H3K27me3 signal in both species. **d,** H3K27me3 overlap profiles for a selection of four GRB TADs. **e,** H3K27me3 overlap profiles of all GRB TADs. TADs are ordered by their relative amount of shared signal. Bins are ordered by the amount of shared signal, bins with shared signal appear in the middle, bins with zebrafish and mouse specific signal are left and right, respectively. A view of the TADs with their genomic bin order is given in **Figure S15d**. **f,** Classification of zebrafish ATAC-seq peaks in the *Irx3a* locus into DC, IC and NC based on overlaps with direct anchors, multi-species anchors and mouse DNase-seq peak projections (see Methods). **g,** Distribution of DNase-seq signal in the mouse genome around the projected regions of the zebrafish ATAC-seq peaks. Asterisks above the bars indicate the effect size category based on cohen’s d: very small (not indicated), small (*), medium (**), large (***), very large (****)^64^. **h,** Cross-species comparison of ChromHMM functional states. **i,** Cumulative distribution of shared motifs in mouse DNase-seq peaks overlapping zebrafish ATAC-seq peaks. **j,** H3K27ac enrichment (signal ≥ 80th percentile) within and outside of enhancer ensembles. **k,** Cross species comparison of H3K27ac profile around an H3K27ac ensemble neighbouring the zebrafish *aktip* gene.

We used this approach to compare zebrafish and mouse TADs, which vary in size approximately two-fold (**Suppl. Figure 15a**). For this purpose, we defined GRB TADs as the 1000 TADs with the highest CNE density, split them into 1kb bins and mapped the bin centers from zebrafish to mouse. Using the multi-species approach over only direct alignments reduced distances from the bin centers to their closest anchor by a factor of 16 and 29 in zebrafish and mouse, respectively (**Figure 6b**).

We then asked whether this method can discover conserved epigenomic subdomains and functional elements with maintained relative locations by comparing epigenomic feature distribution across the genomes. We used H3K27me3 ChIP-seq data from phylotypic stages in zebrafish (Prim-5 stage) and mouse (E10.5 stage; see Methods), when their transcriptomes are most similar^63^. H3K27me3 feature coordinates from zebrafish were projected onto the mouse genome, recovering mouse H3K27me3 features in the corresponding region. An example at the *irx3a* locus (**Figure 6c-e**) shows H3K27me3 enrichment correlates between zebrafish and mouse, even in the absence of direct sequence conservation. This result suggests that epigenomic subdomains and functional elements can be conserved in location and span. We also see more GRB TADs showing regions of strong similarity in H3K27me3 extent, whereas others, such as TADs containing *her9* or *celf5a*, show more mouse or zebrafish specific signal enrichment, respectively, and others show very little enrichment at all (**Figure 6d-e**).

We next looked at conservation of functional elements marked by open chromatin. We classified zebrafish ATAC-seq peaks in the GRB TADs as directly conserved (DC) if they fall in a region of direct sequence alignment with mouse (16,188 elements, 11.5 %), indirectly conserved (IC) if they do not directly align (6137 elements 4.4%) but are alignable through bridging species, and non-conserved (NC) for all other peaks. An example of this is shown for the *irx3a* locus in **Figure 6f**. Notably, DC and IC elements share regulatory features with their matched counterparts in the mouse, including DNase hypersensitivity and ChromHMM feature classification, compared to NC elements (**Figure 6g,h**). DC and IC regions are also more likely to share DNA binding motifs compared to non-overlapping, randomly sampled mouse DNase-seq peaks within and across TAD boundaries (cis and trans in **Figure 6i**; **Suppl. Table 10**). Taken together, these results suggest a similar level of functional conservation of DC and IC elements, even though IC elements lack direct alignability. Next, we tested whether the early developmental H3K27ac ensembles detected in zebrafish embryos (see **Figure 5**) are conserved in the mouse by using our indirect anchoring-based syntenic comparison approach. As shown in **Figure 6j,k**, H3K27ac signal in mouse is enriched in zebrafish ensembles suggesting these ensembles are epigenetic subdomains evolutionarily conserved among vertebrates.

Together, our comparative epigenomic approach has maximised the identification of putative functional elements and epigenetic subdomains conserved between zebrafish and mouse, and highlights the utility of the DANIO-CODE annotations for discovery of vertebrate-conserved mechanisms of developmental gene regulation.

## Discussion

In this study we reported the establishment of a zebrafish developmental genomics resource for the scientific community that consists of comprehensively collected and reprocessed data from both publicly available and newly produced datasets. We describe the provision of this genomics resource as a track hub and as a downloadable resource within a Data Coordination Centre (DCC). The DCC was designed to encourage further expansion by continued incorporation of accumulating zebrafish genomics data. Our track hub provides a reference of developmental non-coding functional annotations and is available for visualisation in common genome browsers and among the Ensembl genomic annotation resources.

We have demonstrated the annotation of over 140,000 candidate developmental cis-regulatory elements including enhancers and promoters. There is a need for the classification of cis-regulatory elements to reflect their distinct modes of functionality or distinct temporal and spatial dynamics. Recognising this need, we improved the functional classifications of both enhancers and promoters and detected previously unknown subcategories by using a novel approach of dimensionality reduction by UMAPs on complex topology data of chromatin accessibility and histone modifications at nucleosome-level. Novel subclasses of cis-regulatory elements include DOPEs and COPEs, possibly described by marks not explored in zebrafish yet^65^, and we demonstrate distinct local chromatin architecture of CREs in developmental stages and in developing cell types. In addition, we classified promoters into novel chromatin architecture-based classes, which we show are distinctly deployed during development by subsets of genes.

We have integrated our CRE annotations with DANIO-CODE chromosome topology data to explore the organisation of CRE interactions during development. We identified large H3K27ac marked domains which we termed H3K27ac ensembles. We show that H3K27ac ensembles are distinct from previously described super-enhancers targeting lineage determining genes, and we assign them to function in nuclear topology organisation at local interaction hubs around early active loci during the initial formation of TADs.

The DCC and the functional annotation track hubs will also serve as a foundational reference for increasing numbers of single cell studies of transcriptomes^66–68^ and that of open chromatin^40^ as was demonstrated with scATAC-seq data in this study. This resource will support integration of tissue and cell-type specificity information with the existing cis regulatory element annotation from bulk data. Future work will allow integration of further layers of functional annotations, including transcription factor binding sites^69^ and integration of TFBS with TF expression data to predict functional binding events relevant for transcriptional activation^70^ or to identify and dissect components of gene regulatory networks acting in lineage determination and organogenesis^71^.

Our high-resolution promoter/ 5’ end transcript annotation will be particularly useful in designing reagents for gene regulation assays^72^, reporter transgenesis for cell labelling^73^, and in transcription blocking experiments (e.g. by dCas9 or Cas9 targeting) where it is necessary to avoid transcription adaptation, which leads to genetic compensation and masked phenotypes in reverse genetic studies of protein coding gene function^45, 74^.

We demonstrate how the utility of our zebrafish functional annotations extends well beyond zebrafish development. We developed a novel approach for the detection of biochemical and biological functional equivalence of regulatory elements and epigenetic landscapes in the absence of sequence conservation with mammals. We exploited conserved synteny together with a novel multistep anchoring by multi-species comparisons, which facilitated direct comparison of non-conserved syntenic locations in fish and mouse TADs with differing lengths. We identified non-sequence conserved positional equivalents with remarkable enrichment for shared features, both on the level of epigenetic domain coverage (e.g. H3K27me3, H3K27ac) and underlying enhancer TFBS content. Our observations expand on the capacity of comparative epigenomics and highlight the predictive value and functional relevance of epigenetic subdomains within syntenic TADs.

This zebrafish resource thus expands on the existing functional genome mapping efforts in mammals and modENCODE species and offers integrated genome annotation resources, with novel observations on promoter and enhancer classes functioning in development and intra-TAD epigenetic feature organisation during development.

## Methods

### Data Collection

We started the DANIO-CODE data collection aiming for capturing a wide range of developing stages in zebrafish from a broad range of genomic, epigenomic and transcriptomic assays.

Members of the zebrafish community were invited to provide their published as well as unpublished data to the DANIO-CODE consortium. Benefiting from experiences of consolidating data in the decentralized data production of the modENCODE consortium^75^, we developed the DANIO-CODE Data Coordination Centre (DCC)^33^ (https://danio-code.zfin.org). The DCC facilitated data collection as well as data annotation and subsequent data distribution.

Demultiplexed FASTQ files were provided by community members to the DCC file system. Using the DCC-web frontend, the community members were guided through an annotation process to annotate the data they provided. The DCC data model is derived from Sequence Read Archive (SRA) data structures^76^ and employs controlled vocabularies based on ZFIN nomenclature^77^. In addition to the community provided data, DANIO-CODE annotators strategically selected additional published datasets to complement developmental stages or assays so far underrepresented in the DCC. These datasets were annotated by the DANIO-CODE curators based on the respective publications.

Consistent data and annotation formats allowed the consistent data processing of all data in the DCC. For this, we developed computational workflows for all the data types and implemented these workflows for the DNAnexus system (https://www.dnanexus.com). Data and annotation quality control measures were established for all data in the DCC.

As a result, all datasets present in the DCC are described in terms of the overall study design, biosamples, library preparation methods, sequencing details as well as in data processing and quality control aspects. Snapshots of the DCC are kept as data freezes to facilitate the handling of newly added data. An interactive data and annotation view and export is provided at https://danio-code.zfin.org/dataExport.

### Data Processing

Sequencing files from the DANIO-CODE DCC were processed with standardised pipelines, see https://gitlab.com/danio-code for details. The resulting files were uploaded to the DCC annotated with the version of the used pipeline and linked to their source files. RNA-seq, methylation data, HiC, 4C-seq data were processed using custom processing pipelines. ChIP-seq and ATAC-seq data were processed using adapted ENCODE pipelines^34^ and CAGE-seq and 3P-seq data were processed using adapted FANTOM pipeline^35^.

### WashU Epigenome Browser tracks

In order to visualize the developmental timeline, bigwig files of the different assays were combined to one matplotlib track and each track was shifted by adding a fixed amount to their score column. To improve readability, the empty regions were set to a score of 0, before the shift. Furthermore, each assay was scaled to reduce noise. The code and the session file are available on GitHub https://github.com/DANIO-CODE/DANIO-CODE_Data_analysis/tree/figure_1/Figures/Figure1. Zebrafish developmental stage drawings were adapted from Kimmel et al., 1995^78^.

The drawing of the adult stage in **Figure 1** was taken from https://www.drawingtutorials101.com/how-to-draw-a-zebrafish.

### Nanti-CAGE-seq on zebrafish developmental stages

Total RNA was extracted from multiple stages of zebrafish development (Fertilized egg, 16-cell, 128-cell, 512-cell, 30%-epiboly, 4 somite, Prim-5, and High-pec) using the miRNeasy kit (Qiagen), according to the manufacturer’s instructions. nAnT-iCAGE libraries were prepared as described previously^79^ using the CAGE™ Preparation Kit (DNAFORM). All libraries were sequenced on Illumina HiSeq2500 except the high pec library which has been sequenced on NextSeq500.

### Total and Nuclear Tagging CAGE-seq

Total RNA was collected from multiple stages of zebrafish development (30%-epiboly, 9 somite, Prim-5, High-pec and Larvae) using the miRNeasy kit (Qiagen), according to the manufacturer’s instructions. Nuclear RNA was also collected, through a nuclear isolation protocol based on the Nuclei EZ isolation kit (Sigma) but optimized for zebrafish tissue. In summary embryos were de-corionated and then dissociated and de-yolked in a cell swelling buffer (250mM Sucrose, 10mM Tris-HCL (pH 7.9), 10mM MgCl_2_, 1mM EGTA), through vigorous pipetting. Embryo slurry was left to stand for 5 minuted and then filtered using a 50uM tube top filter. It was then centrifuged at 500g, 5 mins, 4°C and the pellet resuspended in Freezing buffer (Sigma), after which they were stored at −80°C. Upon defrosting Nuclei EZ Lysis Buffer was added to the cells, vortexed and left to stand for 5 mins. Nuclei were then centrifuged at 500g, 5 mins, 4°C and the pellet washed once in Nuclei EZ Lysis Buffer. Upon a final centrifugation of the nuclei, the pellet was resuspended in Qiazol Lysis Reagent (Qiagen) and RNA extracted using the miRNeasy kit (Qiagen), according to the manufacturer’s instructions. Tagging-CAGE libraries were prepared from both the total and nuclear fractions following the protocol described previously^80^. All libraries were sequenced on Illumina HiSeq2500 by the SNP&SEQ Technology Platform in Uppsala, Sweden.

### Embryo preparation and ChIP experiments

Approximately 1500 embryos Dome/30%-epiboly, 25 embryos 5-9 somites and 200 embryos Prim-5/Long-pec were collected. Embryos were enzymatically dechorionated using pronase and fixed in 1.85% formaldehyde in Hanks media for 20 minutes at room temperature. After crosslinking, samples were washed once with PBS and the fixation was stopped by incubating in 1x glycine for 10 minutes at room temperature followed by three washes with ice-cold PBS.

The embryos at somite stages were fixed in 1% formaldehyde for 10 minutes at room temperature upon dissociation in 1 ml of PBS and 1x protease inhibitor cocktail. Fixation was stopped with 115 μl of glycine for 5 minutes at room temperature. The cell suspension was centrifuged at 300x g for 10 minutes at 4°C and pellets were washed with Hank’s balanced salt solution (HBSS) supplemented with 1x protease inhibitor cocktail (PIC) and centrifuged at 300x g for 10 minutes. Supernatants were removed and pellets were kept on ice.

ChIP experiments were carried out using the ChIP-IT Express Enzymatic kit (Cat. No. 53009, Active Motif) or the True MicroChIP Kit (Cat. No. C01010130, Diagenode) for the 5-9 somite stage embryos. Both protocols were applied in line with manufacturer’s instructions.

Briefly, for the ChIP-IT Express Enzymatic Kit, crosslinked embryos were resuspended in 1 ml ice-cold lysis buffer, incubated on ice for 20 minutes, transferred to a pre-cooled dounce homogenizer and dounced by 10 strokes. Nuclei were collected by centrifugation, resuspended in 200 μl digestion buffer and incubated at 37°C for 5 minutes. Chromatin was sheared by adding 10 μl of enzymatic shearing cocktail working stock (200 U ml−1) and incubating for 10 minutes at 37°C. Shearing efficiency was checked by gel electrophoresis according to manufacturer’s instructions. The reaction was stopped by adding 5 μl ice-cold 0.5 M EDTA and incubating on ice for 10 min and sheared chromatin was cleared by centrifugation. For ChIP reactions 70 μl of sheared chromatin were mixed with 25 μl Protein G magnetic beads, 20 μl ChIP buffer 1, 1 μl protein inhibitor cocktail and 4 μg of antibody or an equivalent volume of water (no antibody control), respectively, and water to a final volume of 200 μl. ChIP was performed in duplicates for each stage. ChIP reactions were then incubated in rotation overnight at 4°C. Magnetic beads were washed and incubated in elution buffer. After addition of reverse crosslinking buffer samples were decrosslinked for 4 hours at 65°C. Samples were treated with Proteinase K and RNase A and purified using phenol chloroform extraction.

ChIPs for 5-9 somite stage embryos were performed with the True MicroChip in crosslinked and dissociated embryos as described above. Cell pellets were incubated with 25 μl lysis buffer and 1x PIC for 10 minutes on ice. Upon addition of 75 μl of HBSS the cell suspension was sonicated with 12 cycles (30 seconds ON/ 30 seconds OFF) using a Bioruptor Pico sonicator. Sonicated samples were centrifuged at 14000 x g for 10 minutes and the supernatant was mixed with 100 μl of tC1 buffer and 1x PIC. Chromatin was incubated with the antibody in rotation overnight at 4°C together with 11 μl of pre-washed. Chromatin-bound beads were captured using a magnetic rack and washed with 100 μl tW1, tW2, tW3 and tW4 buffer respectively, for 4 minutes at 4C each time. Beads were resuspended in elution buffer tE1 and incubated for 30 minutes at room temperature. Finally, supernatants were transferred to clean tubes and de-crosslinked with 8 μl of Elution buffer tE2 for 4 hours at 65C. DNA was purified using Micro ChIP spin columns.

#### Antibodies used for ChIP-seq

| Mark | Reference |
| --- | --- |
| H3K4me3 | C15410003 (Diagenode) |
| H3K27ac | ab4729 (abcam) |
| H3K4me1 | ab8895 (abcam) |
| H3K27me3 | C15410069 (Diagenode) |
| H3K36me3 | ab9050 (abcam) |
| Pol II | ab5131 (abcam) |
| H2A.Z | ab4174 (abcam) |

### Library preparation for ChIP-seq experiments

The DNA libraries were prepared using the Microplex library preparation Kit V2 (Diagenode, Cat. No. C05010013) following the manufacturer’s instructions. Briefly, templates were prepared with 2 μl of template preparation buffer and 1 μl of preparation enzyme and the reaction was carried in a thermocycler as follows: 25 minutes at 22°C, 20 minutes at 55°C and hold at 4°C. Upon template preparation libraries were synthesized with 1 μl of library synthesis buffer and 1μl of library synthesis enzyme incubate the samples at 22°C for 40 minutes before proceeding to library amplification. Finally, libraries were amplified adjusting the number of cycles to the amount of DNA used for the preparation, according to the instructions. All libraries were purified and size selected with AMPure XP beads (Beckman Coulter) using the ratios recommended by the microplex protocol. The quality and concentration of the samples was verified by High Sensitivity D5000 ScreenTape (Agilent) and Qubit dsDNA High sensitivity assays (Thermo fisher scientific).

### ATAC-seq experiments

Chromatin was prepared using 200 embryos, 100 embryos and 50 embryos in the 32-cell, 128-cell and 256-cell stages, respectively. Embryos were dechorionated enzymatically using pronase and collected in 1.5ml eppendorf tubes. 1ml Cell Dissociation Buffer (13151014, Gibco) was added into the tubes and cells were dissociated by pipetting up and down for several times. Dissociated cells were then collected by spinning at 500g for 10min at 4°C. The following steps followed the ATAC protocol as previously published^47^.

### Cell preparation for Hi-C

Samples were prepared as previously described^81, 82^. Briefly, Tg(Buc-GFP) heterozygous embryos were dissociated in 500 ml of HBSS supplemented with 0.25% BSA and 10mM Hepes by pipetting for 2 minutes with a glass pipette. Excess of yolk was removed by two rounds of 3 minutes centrifugation at 350 x g at 4°C, while pelleted cells were resuspended in 1ml HBSS supplemented with 0.25% BSA and 10mM Hepes prior to filtering. The cell suspension was kept on ice during sorting in a FACS Aria Fusion sorter.

### Low input somatic cells in situ Hi-C library generation

Briefly, we performed low input in situ Hi-C^82^ with small modifications on sorted somatic cells in biological duplicates. 6,000 FACS-sorted somatic cells were crosslinked in 1% formaldehyde, incubated for 10 min at room temperature with rotation (20 rpm) and quenched by adding 0.2M glycine for 5 min at room temperature with gentle rotation (20 rpm). Cells were then washed three times with 1 mL cold PBS (centrifuged at 300 g for 5 min at 4C) and lysed using ice-cold in situ Hi-C buffer (10 mM Tris-Cl pH 8.0, 10 mM NaCl, 0.2% IGEPAL CA-630, cOmplete Ultra protease inhibitors) by incubating them on ice for 15 min. Centrifuged nuclei were then resuspended in 125 mL ice-cold NEB2 buffer, pelleted (13,000 g for 5 min at 4C) and permeabilized inn12.5 mL of 0.4% SDS at 65C for 10 min. SDS was quenched by adding 12.5 mL of 10% Triton X-100 and 85 mL of nuclease-free water and incubating at 37C for 45 min with shaking. Chromatin was digested with 100 U of MboI restriction enzyme (90 min at 37C with rotation; New England Biolabs) and the enzyme was heat-inactivated at 62C for 20 min. Digestion generated overhangs were filled in with a mixture of 0.4mM biotin-14-dCTP (10 mL; Thermo Fisher Scientific) and 10mM dATP/dGTP/DTTP (0.5 mL each; Thermo Fisher Scientific) by incubating with DNA polymerase I Klenow (4 mL; New England Biolabs) for 90 min at 37C with rotation. DNA fragments were ligated using T4 DNA ligase (Thermo Fisher Scientific) and a mix (60 mL of T4 DNA ligase buffer, 50 mL of Triton X-100, 6 mL of BSA 20mg/ml, 3.5 mL of T4 DNA ligase and 328.5 mL of nuclease-free water) during 4 h at 20C with gentle rotation. Nuclei were gently pelleted (2,500 g for 5 min at room temperature) and resuspended in 250 mL extraction buffer. Protein was digested with 10 mL Proteinase K (20mg/ml; Applichem) for 30 min at 55 C with shaking (1,000 rpm) followed by adding 65 mL of 5M NaCl and an overnight incubation at 65C with shaking (1,000 rpm). Phenol-Chloroform-Isoamyl alcohol (25:24:1; Sigma-Aldrich) DNA extraction was performed on the nuclei and final DNA was resuspended in 25 mL of 10 mM Tris pH 8.0 (Applichem) and incubated with RNase A (1 mL of a 10mg/mL stock; Applichem) for 15 min at 37C. Biotin removal from unligated fragments was performed by incubating DNA samples for 4h at 20C in a mix of 5 mL of 10X NEB2 buffer (New England Biolabs), 5 mL of a 1mM dNTPs mix, 0.25 mL of 20 mg/ml BSA, 3.5 mL of T4 DNA polymerase (New England Biolabs) and up to 50 mL of nuclease-free water. Samples were then sheared using a Covaris S220 instrument (2 cycles, each 50s, 10% duty, 4 intensity, 200 cycles/burst). Dynabeads MyOne Streptavidin C1 beads were used to pull down biotinylated fragments. NEBNext Ultra End Repair and the NEBNExt Ultra dA-Tailing modules were used for end repair and adding Illumina sequencing adaptors. Final PCR amplification was done in 4 parallel reactions per sample (10 mL of the bead-bound libraries, 25 mL of 2x NEBNext Ultra II Q5 Master Mix, 3 mL of 10 mM Universal PCR primer from Illumina, 3 mL of 10mM Index PCR primer and 9 mL of nuclease-free water and amplified for 10 cycles (98C for 10 s, 65C for 75s, ramping 1.5C/s for 10 cycles and a final step at 65C for 5 min). The four reactions were combined into one tube and size selected using Ampure XP beads (Beckman Coulter). Final Hi-C libraries were quantified using Qubit dsDNA HS assay kit and a DNA HS kit on a 2100 Bioanalyzer (Agilent). Libraries were first shallow sequenced on an Illumina MiSeq (2×84bp paired-end; MiSeq reagent kit v3-150 cycles) to assess library quality. Finally, libraries were deeply sequenced on an Illumina NextSeq (2×80 bp paired-end; NextSeq 500/550 High Ouput kit v2-150 cycles).

### 4-C data generation

4C-seq experiments were performed and analysed as described earlier^83^. For zebrafish, 500 embryos at the 24hpf stage were dechorionated with pronase and deyolked using the Ginzburg Fish Ringer buffer (55 mM NaCl, 1.8 mM KCl and 1.25 mM NaHCO3). Then they were fixed using 2% PFA for 15 minutes at room temperature. Fixations were stopped by adding glycine and washing several times with PBS. Then, the fixed samples were lysed (lysis buffer: 10 mM Tris-HCl pH 8, 10 mM NaCl, 0.3% Igepal CA-630 (Sigma-Aldrich, I8896) and 1× protease inhibitor cocktail (Complete, Roche, 11697498001)), and the DNA was digested with DpnII (New England BioLabs, R0543M) and Csp6I (Fermentas, Thermo Scientific, FD0214) as primary and secondary enzymes, respectively. T4 DNA ligase (Thermo scientific, EL0014) was used for both ligation steps. Specific 4C-seq primers containing single-end Illumina adaptors were designed using as bait the different promoters of interest. For each library, eight independent PCRs were performed using the Expand Long Template PCR System (Roche, 11759060001) that were subsequently pooled and purified using AMPure XP beads (Beckman Coulter, A63883). Then libraries were sent for single-end sequencing. 4C-seq sequencing reads were aligned to the zebrafish reference genome using bowtie. Reads located in fragments flanked by two restriction sites of the same enzyme, in fragments smaller than 40 bp or within a window of 10 kb around the viewpoint were filtered out. Mapped reads were then converted to reads per first enzyme fragment ends and smoothened using a 30-fragment mean running window algorithm. This signal is ready to be visualized. More details and supporting code are available in the GitLab repository (https://gitlab.com/danio-code/danio-code_4c-seq).

### Promoterome construction

First, all reads mapping to poorly assembled genomic regions or otherwise blacklisted^84^ regions were excluded from the set of cTSSs. After an initial application of CAGEr^85^ we discovered systematic differences between nAnTi and tagging CAGE samples both at the number of Transcription Start Site Clusters (TCs) number and summed promoter/gene expression across samples produced with the two CAGE protocols. Particularly, the fraction of CAGE signal coming from annotated exons was elevated in nAnTi samples skewing the statistics. To a varying extent this phenomenon (known as exon painting/carpeting) has been previously observed and attributed to recapping of degraded mRNAs. Since the majority of true TSS are initiated at either YC or YR dinucleotides^13^, we analysed dinucleotide frequencies at initiation sites and confirmed an increased proportion of other (non-YC and non-YR) dinucleotides in nAnTi compared to tagging samples. We therefore decided to remove all CAGE tags not initiated at YC or YR dinucleotides.

The remaining set was of TSSs was power-law normalised^86^ to a common exponent alpha=1.1 and 5 to 1000 tags fit range, and the TCs were produced using the following parameters of CAGEr: *threshold = 0.7, thresholdIsTpm = TRUE, nrPassThreshold = 1, method = “distclu”, maxDist = 20, removeSingletons = TRUE, keepSingletonsAbove = 5*

This yielded a comparable number of TCs across samples without an obvious bias towards high numbers in nAnTi samples. The number of TCs moderately increased in post-ZGA samples.

### Number of TCs per sample

To compare expression levels across samples we called consensus clusters (genomic regions not assigned to any particular sample unlike TCs) with settings *tpmThreshold = 1.0, qLow = NULL, qUp = NULL, maxDist = 20*. To further filter weak or spurious tag clusters we kept consensus clusters which were expressed in at least 2 consecutive developmental stages. Specifically, we required that there exist a TCs within a consensus cluster in both consecutive stages with at least 1.0 tpm expression. This yielded 27781 consensus cluster. We calculated expression of each consensus cluster by summing all YC and YR-initiated tag CAGE tags from within the cluster across stages. This approach differs from CAGEr implementation which includes expression only from TCs within a consensus cluster and is subject to generating noise at lowly expressed regions due to *tpmThreshold* parameter.

To visualise in such way obtained expression levels we made a 2D PCA plot, which correctly grouped nAnTi CAGE and tagging CAGE samples from the same stage.

### Expression clustering

We clustered 27781 consensus cluster expression log fold changes with self-organising maps choosing a 5 x 5 grid as in previous work^87^. These analyses separated consensus clusters mostly by maternal and zygotic expression. Afterwards we manually grouped the resulting 25 cells into maternal (blue), zygotic (red), ZGA-peaking (brown) and constitutively expressed at different levels (gray).

### Promoter classification by open chromatin

For each consensus cluster we defined a reference point as the TSS with highest mean post-MBT expression “dominant TSS” (tpm values of samples ranging from the Shield to Long-pec stages) and required that it amounts to at least 0.2 tpm. This further reduced the set of consensus clusters to 21914 elements. We then merged ATAC samples from the Prim5 stage and extracted Tn5 cut sites from both ends of ATAC reads while correcting for Tn5 overhang, smoothed with a Gaussian kernel with standard deviation 3bp and log-transformed. These ATAC cut-site profiles served as input to k-means clustering (k=8, range +-800bp from the dominant TSS).

### Transcripts identification

Wild-type embryonic paired-end and stranded RNA-seq samples (DCD000141SR, DCD000225SR, DCD000247SR, DCD000433SR, DCD000426SR, DCD000324SR)^16, 41, 43, 88–90^ were selected from a total of 528 DANIO-CODE RNA-Seq and aligned to GRCz10 using STAR aligner v. 2.5.1b^91^. StringTie v.1.33b^92^ was used to call transcripts, which were then all assembled using TACO^93^, generating a total of 194,508 transcripts in canonical chromosomes. Transcript quantification was done using Salmon v.0.11.2^94^. We removed read through, mono-exonic and transcripts overlapping 3 or more Ensembl genes. All protein coding transcripts above 200kb and long non-coding RNAs above 100kb were excluded. We selected transcripts that are expressed in minimum 2 closest stages out of the remaining set (permissive set, n=66,341), yielding 51,033 transcripts (robust set).

### Guide RNA preparation and microinjection

Single guide RNAs (dried pellet, Agilent) were resuspended with H2O into 100mM stock concentration. Zebrafish codon optimized dCas9-Neon was cloned into pCS2+ vector, then the plasmid was linearized with NotI and dCas9-Neon mRNA was synthesized *in vitro* with mMESSAGE mMACHINE™ SP6 Transcription Kit (Ambion). Single guide RNA (80nM/ul) and dCas9neon mRNA (500ng/ul) were injected into newly fertilized one cell stage zebrafish embryos at 2nl volume.

### Annotation of alternative transcript and alternative promoter

A gene can have multiple transcripts/isoforms that differ in their transcription start sites by few nucleotides to tens of kilobases. When a gene has multiple transcripts, Ensembl assigns the longest transcript as a reference transcript for that particular gene and its promoter is defined as reference promoter. To determine non redundant alternative promoters, we analyzed annotated transcripts models from Ensembl and novel RNA-seq transcripts models from DANIO-CODE. Similar to that of the Ensembl method, we annotated the longest transcript as the reference transcript and its promoter as reference promoter. Remaining transcripts whose transcription start sites were proximal (<300 nucleotides) to reference transcripts were excluded and remaining distal transcripts were used to determine alternative transcripts/promoters. The longest transcript was assigned as an alternative transcript and excluded other transcripts with proximal (<300 nucleotides) transcription start sites. As some genes have more than one alternative transcripts, we iterated this process to annotate additional alternative promoters that are at least 300 nucleotides apart from already assigned reference or alternative transcripts.

To identify conserved alternative transcripts/promoters in humans, we used human/zebrafish ortholog table from Ensembl^95^ and annotated gene/transcript models from Ensembl annotation. We first intersected 1253 zebrafish genes with alternative promoters and identified 1055 genes that have human orthologs. Then we intersected 1055 human orthologous genes with Ensembl transcript models and identified 929 genes that also had annotated alternative transcripts in humans.

### Genome browser visualisation of chromatin marks and annotated regions

Chromatin marks and gene tracks (Figure 4A, Figure 5D), as well as the corresponding annotation tracks (ChromHMM, PADREs, and ensembles) were visualized with the Gviz package^96^.

### Functional segmentation of the genome

In addition to well-annotated genes, the genome contains tens of thousands of regulatory elements regulating gene expression. Identification of such regulatory elements is challenging. Luckily, to be active, regulatory elements must modify their histone proteins with characteristic covalent modifications. For example, acetylation of lysine residue 27 and mono-methylation of lysine residue 4 on the histone protein H3 (H3K27ac and H3K4me1) are a feature of active chromatin. The modification H3K4me3 is characteristically found on promoters, while H3K27me3 represents Polycomb repressed chromatin. Those specific modifications can be localized using methods such as chromatin immunoprecipitation followed by sequencing (ChIP-seq). Algorithms such as ChromHMM^48, 49^ can segment the genome into regions containing specific chromatin marks. Using the chromatin marks mentioned above, we segmented the genome capturing the epigenetic state of the genome in 5 different development stages. We used published data for the Dome, 75%-epiboly, 5-9 somites, Prim-5, and Long-pec stages^9, 41, 87, 97–99^, as well as newly produced data for the 5-9 somites and Long-pec stages. After optimization, we found ten optimal latent states based on the emission parameters of chromatin marks. The states were matched between stages and manually assigned a function using The Roadmap Epigenome Project^28^ annotation as a reference. The identified functional elements were annotated as followed:

1. Active TSS 1 (1_TssA1),
2. Active TSS 2 (2_TssA2),
3. TSS Flanking region 1 (3_TssFlank1),
4. TSS Flanking region 2 (4_TssFlank2),
5. Active enhancer 1 (5_EnhA1),
6. Weak enhancers (6_EnhFlank),
7. Primed enhancer (7_EnhWk1),
8. Poised elements (8_Poised),
9. Polycomb repressed regions (9_PcRep),
10. Quiescent state (10_Quies)

The active promoters and promoter flanking regions, in addition to active chromatin marks, show strong emission of H3K4me3. Moreover, the promoter-associated states are mostly found on and around the annotated TSS (Figure 4a, middle and right). The states missing H3K4me3 and not found around TSS were annotated as enhancer-related. Depending on if both H3K27ac and H3K4me1 were present, as well as the strength of the emission, the enhancer states were divided into active enhancers (strong H3K27ac and H3K4me1 emission, but no H3K4me3), enhancers flanking (weak H3K27ac emission, mostly found around active enhancers), and primed enhancers (H3K4me1 emission only). States emitting H3K27me3 were annotated as Polycomb-related. In addition to H3K27me3, when active marks were present, the state was assigned as poised; otherwise, it was assigned as Polycomb repressed. When no marks were present, the region was assigned as quiescent. Most of the genome shows no marks at all.

### Predicted ATAC-supported Developmental Elements (PADRE)

Although the ChromHMM segmentation points to a presence or absence of a mark, it does not tell about the actual activity of the elements. Moreover, it has a 200bp resolution, and it does not consider the peak nor the profile of the signal. We constrained our subsequent analyses to the regions in the genomes which are open, i.e., depleted in nucleosomes as identified by ATAC-seq. We identified stage-specific open chromatin regions consistent between replicates with the irreproducibility discovery rate^100^ less than 0.1 in seven developmental stages (4 pre-ZGA stages, newly produced datasets, and 7 post-ZGA stages of which 30%-epiboly is newly produced and the other samples were published previously^90^). We termed those regions as predicted ATAC-supported developmental regulatory elements (PADRE). All stage-specific PADREs were merged to form a set of regions called consensus PADREs (cPADREs). Two different sets of cPDREs were defined. The permissive cPDREs comprise all PADREs merged and number around ∼240k elements. The strict set takes into account only regions that are open in at least two neighbouring stages. This set counts ∼140k elements. All cPADREs analysis in this paper were done on strict cPADREs.

Although ATAC-seq can very precisely and robustly detect open chromatin regions pointing to their activity, it does not give much information regarding the function of the identified elements. To address this question, we assigned each ATAC peak a ChromHMM state based on overlaps. Overall, the total number of promoter-associated regions is comparable across the stages, with minor discrepancies between in-group states.

### DNA Methylation analysis

Bisulfite-converted (WGBS) sequence reads from developing zebrafish embryos^18–20^ were trimmed using Trimmomatic with the following settings: ILLUMINACLIP:TruSeq3-SE.fa:2:30:10 SLIDINGWINDOW:5:20 LEADING:3 TRAILING:3 MINLEN:20^101^. Using Walt (-m 5 -t 20 -N 10000000)^102^, trimmed reads were mapped onto bisulfite-converted reference genomes for both GRCz11 and GRCz10 (UCSC) with the λ genome added as a separate chromosome in order to estimate bisulfite conversion efficiencies. The resulting SAM files were converted to BAM format and were deduplicated using sambamba markdup^103^. Due to some samples containing high levels of non-conversion rates (>1%), BAM files were filtered to remove reads containing more than 3 non-converted cytosines outside the CG context using samtools^104^, picard (http://broadinstitute.github.io/picard/) and a custom awk script. Methylation levels were called using MethylDackel (extract --mergeContext) (https://github.com/dpryan79/MethylDackel), and Bigwig files were generated from the resulting bedGraph files using UCSC bedGraphToBigwig^105^. Matrices for heatmaps were generated using Bigwig files and deepTools^106^ (computeMatrix reference-point -b 2500 -a 2500 -bs 25). NAN values from the resulting Matrix files were replaced with zeroes and heatmaps were plotted using deepTools plotHeatmap. Average methylation levels for chromHMM analyses were calculated using BEDtools^107^ (map -o sum) by dividing the sum of reads supporting a methylated CG by the number of reads mapped to that region.

### UMAP visualisation

We developed a method that considers various signals around the open chromatin summit to address all those concerns. In brief, we constructed a feature matrix using ATAC-seq, H3K4me3, H3K27ac, H3K4me1 ChIP-seq tags, as well as nucleosome position calculated by NucleoATAC^108^. Nucleosome signal was included because some factors have well-positioned nucleosomes around their binding sites and could separate those factors from others. In brief, the peak summit is extended for 750bp in each direction and split into 13 bins (R1 - R13). For each bin, the number of tags for the aforementioned assay types is counted, and the mean nucleosome signal in each bin was calculated using the genomation package. It resulted in five score matrices, each having the number of rows the same as the number of open chromatin regions and 13 columns (one for each bin around the peak summit). Those matrices were standardised by scaling the values and centring the mean to 0. The standardised matrices were concatenated column-wise, giving a total of 65 columns (**Suppl. Figure 8a-c**). Using the UMAP algorithm^109^, the number of features was reduced from 65 to 2, making it possible to plot each open chromatin region into a 2D plot.

### eRNA calling

We identified eRNA using CageFightR v1.8.0^110^ on the 24 total and nuclear Tagging CAGE-seq samples with the standard parameters. We applied an expression filter, keeping only clusters with an expression greater than zero (`unexpressed=0`) in more than two samples (`minSamples=2`). Furthermore, only intergenic and intronic clusters were kept. The code is available in the GitHub repository.

### Identification of CTCF binding sites

The CTCF PWM matrix was taken from the JASPAR database^69^ under the ID MA0139.1. PADREs were scanned for CTCF motif using TFBSTools^111^, using a threshold of 90% identity.

### Conserved Non-Coding Elements (CNEs) and phastCons

CNEs used were downloaded from Ancora (http://ancora.genereg.net)^112^. Human CNEs used were 70% matched over 50 columns, and mexican cavefish CNEs used were 90% matched over 50 columns. Cyprinid phastCons score^113^ (grass carp, common carp, goldfish, adn zebrafish) for the genome version danRer10 were downloaded from https://research.nhgri.nih.gov/manuscripts/Burgess/zebrafish/download.shtml^114^.

### Cell-type specificity assignment

Genome-wide cell-type specificity of the genome version of danRer11 was assigned from single-cell ATAC-seq data as in McGravey et al., 2020. PADREs were converted from the danRer10 version to danRer11 using the *liftOver* function from the *rtracklayer* package^115^. The chain file for lift-over was downloaded from the UCSC genome browser^36^. The ATAC-seq, H3K27ac ChIP-seq, and H3K4me1 ChIP-seq signals of brain, muscle, and differentiating neuron specific elements were visualized with the *genomation* package^116^ with a window of +/-750bp around the peak summit.

### Classifications of distal elements by chromatin opening dynamics throughout development

Distal cPADREs were filtered by excluding cPADREs +/- 500bp around CAGE-defined promoters. For each distal cPADRE the fold change of the ATAC-seq signal over background was calculated at every developmental stage. Each distal cPADRE were normalized stage-wise by dividing it’s value by root-mean-square and those values were used as input for the self-organizing maps (SOMs)^117^. SOMs were performed using the *kohonen* package^118^. The clustering was performed using a 4×4 hexagonal grid, which resulted in 16 classes ordered in a 4×4 grid. The ATAC-seq, H3K27ac ChIP-seq, and H3K4me1 ChIP-seq signals of elements assigned to classes 4, 6, and 14 were visualized with the *genomation* package^116^ with a window of +/-2000bp around the peak summit. Peak summits for each stage were defined with the *refinepeak* command from the MACS2 program package^119^ using the Dome and Prim-5 stage-specific mapped reads.

### Constitutive Orphan Predicted Elements (COPEs) and Dynamic Orphan Predicted Elements (DOPEs)

Constitutive elements were defined as the intersection of distal PADREs at every developmental stage. COPEs were defined as constitutives annotated as quiescent at every developmental stage. DOPEs were defined as cPADREs, annotated as quiescent at every developmental stage. DOPEs were further classified as adults-marked DOPEs if they overlapped H3K27ac marked regions in any of the adult tissues^31^.

### Identification of topologically associated domains (TADs)

Long-pec (48 hpf) Hi-C matrices (Hernandez-Rodriguez, Díaz, et al., Locus-specific functional conservation of chromatin structure across vertebrate evolution. In revision) were mapped to the GRCz10 genome and processed using FAN-C v0.9.0 with default parameters^120^. Merged matrices were binned at 10kb resolution. TADs were interactively called using TADtool^121^ with a window size of 110kb and a cut-off of 0.004148 over the insulation index signal.

### Identification of Genomic Regulatory Blocks (GRBs)

The GRBs used were called from whole genome alignment between zebrafish and mexican cavefish as described previously^122^, with the following parameters: 70% match and 18kb window. Extreme conservation is a (fat) tail of a continuum, so some “non-GRB TADs” might be TADs with higher turnover of noncoding conservation. Our comparison looked for enrichments, not sharp differences.

### H3K27ac ensemble identification

Enhancer ensembles were detected using H3K27ac peaks and mapped reads from the Dome stage (DCC Data IDs: DCD006167DT and DCD008973DT) as input for the ROSE algorithm^57^ with the distance from TSS to exclude, adjusted to 500bp.

### Hi-C data processing and normalisation

Raw sequencing reads were processed using HiCUP v0.6.0^123^. Sequencing reads were mapped against the danRer10 genome, with Bowtie2 v2.3.4.1^123^ as the aligner. Experimental artifacts, such as circularised fragments or re-ligations, were filtered out, and duplicate reads were removed. The aligned Hi-C data were normalised (coverage-corrected) using a matrix balancing algorithm by HOMER v4.11^124^.

### Aggregate contact map analysis around H3K27ac ensembles

Ensembles were categorised as GRB or non-GRB ensembles, and 50-150 kb long ensembles were selected for this analysis. Control sets of ensembles were generated by shifting the ensemble coordinates by 10Mb, or randomly uniformly sampling GRB/non-GRB TADs with/without ensembles. All sets were downsampled to the smallest set. The aggregate coverage- and distance-corrected Hi-C maps around these ensemble sets were computed at 10 kb resolution in a +/-500 kb or +/-150 kb window around the centre of ensembles, using HOMER’s *analyzeHiC* function with the options *-hist 10000 - superRes 10000 -norm -normTotal given -normArea given*.

### Compartment analysis

The compartment signal was computed as the first principal component of the normalised (coverage- and distance-corrected) interaction profile correlation matrix at 50 kb bin resolution using the *runHiCpca.pl* function of HOMER. Chromosomes where the contact enrichment within chromosome arms was stronger than the compartment signal, we used the second principal component. Chromosomes 4 and 7 were excluded from the analysis as the compartment calls were poor quality due to assembly issues. TADs were categorised into 4 categories, based on whether they were GRB or non-GRB TADS and whether they did or did not contain ensembles. For all TADs, a compartment score was defined as the compartment signal of the central 50kb bin of the TAD. The distributions of compartment scores for the GRB/non-GRB TADs with/without ensembles were compared using two-sided two-sample unpaired Wilcoxon test.

### Expression profiles of genes in TADs with H3K27ac ensembles

CAGE promoters located in TADS with H3K27ac ensembles are classified as H3K27ac associates if their promoter is located within the ensemble or in the 12.5kb flanking region of the ensemble. The expression of each promoter in H3K27ac TADs throughout development was centered and scaled, and the promoters were clustered using the *kohonen* package^118^. The clustering was performed using a 3×3 hexagonal grid, which resulted in 9 classes ordered in a 3×3 grid. The obtained classes were visualized using the *ComplexHeatmap* package^125^.

### Expression of target and bystander genes

The list with prediction of human target and bystander genes was used from Tan, 2017. Zebrafish orthologs were identified using a homology table from *biomaRt*^126, 127^. To account for ohnologs, only predicted target genes with a homolog in a GRB TAD were considered in the analysis. The expression of each promoter in GRB TADs throughout development was centered and scaled, and the expression was visualized using the ComplexHeatmap package.

### H3K27ac signal across TADs

H3K27ac signal across TADs was calculated using the ScoreMatrixBin function from the genomation package with 400 bins. Signals of H3K27ac for each stage were used as targets. For windows, each TADs were ordered by length and each TAD was extended to the size of the largest TAD. The calculated matrices were smoothed and visualized using the heatmaps package (Perry, 2021, doi: 10.18129/B9.bioc.heatmaps).

### Track hubs

Processed data track can be uploaded to UCSC Genome browser as a track hub^126^. The DCC track hub was generated using custom scripts and it is available in the following link: https://danio-code.zfin.org/trackhub/DANIO-CODE.hub.txt, but also registered at UCSC as a public track hub. Annotation data are available in the track hub available in the following link: http://trackhub.genereg.net/DANIO-CODE_Functional_annotation_tracks/hub_test/hub1.txt

### Genomic coordinate projection

Genomic coordinates of GRB loci were projected between zebrafish and mouse using multiple pairwise sequence alignments between a set of 15 species. The basic concept of our approach is that - under the assumption of conserved synteny - a non-alignable genomic region can be projected from one species to another by interpolating its relative position between two alignable anchor points. The accuracy of such interpolations correlates with the distance to an anchor point. Therefore, projections between species with large evolutionary distances such as zebrafish and mouse tend to be inaccurate due to a low anchor point density. Including so-called *bridging species* may increase the anchor point density and thus improve projection accuracy. **Suppl. Figure 15b** illustrates the potential benefit of using a bridging species with a schematic example projection between zebrafish and mouse. The optimal choice of bridging species may vary between different genomic locations and there may be genomic locations for which a combination of bridging species with intermediate projections produces optimal results. **Suppl. Figure 15c** presents the bridging species optimization problem as a shortest path problem in a graph where every node is a species and the weighted edges between them represent the distance of a genomic location to its anchor point. For that, we established a scoring function that reflects those distances and returns values between 0 and 1, where a score of 1 means that a genomic location x overlaps an anchor point a. The score decreases exponentially as the distance |x − a| increases. For a single species comparison, the function is defined as follows:

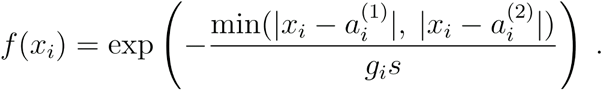

with g denoting the genome size of the respective species and s a scaling factor that can be determined by defining a *distance half life* d_h_, as the distance |x-a| at which the scoring function is to return a value of 0.5:

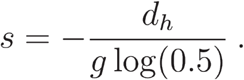

The length of a path through the graph is then given by subtracting the product of the distance scoring function for every node in the path from 1:

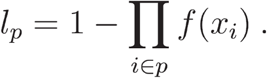

The shortest path *p̂* through the graph is then found by minimizing l_p_:

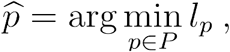

with P denoting the set of all paths through the graph. The optimization problem presents a classic shortest path problem and is solved using Dijkstra’s Shortest Path Algorithm^128^.

### Epigenomic Profile Comparison

We compared H3K27me3 ChIP-seq data from phylotypic stages in zebrafish (Prim-5 stage) and mouse (E10.5 stage)^129^, when their transcriptomes are most similar^63^. In order to match the whole-embryo zebrafish data, we created a virtual embryo dataset for mouse by merging data for 6 different tissues (fore-, mid-, hindbrain, facial prominence, heart, limb). The mouse H3K27me3 profile was projected onto zebrafish genomic coordinates using the multi-species approach by splitting the zebrafish GRB into 1 kb windows, projecting their center coordinates onto mouse and retrieving the signal from the respective 1 kb bin in mouse. ‘Signal’ stands for H3K27me3 coverage represented as quantiles after quantile normalization of the two distributions in zebrafish and mouse. Signal overlap is represented by the log signal ratio and capped to values in [-1,1]. Signal amplitude represents the maximum signal of zebrafish and mouse to the power of 10 in order to increase the variance of signal amplitude. For mouse and zebrafish comparison of H3K27ac ensembles previously published data were used^130^.

### Classification of Conservation

Zebrafish ATAC-seq peaks were classified into three conservation classes based on the projection using the multi-species approach. Directly conserved (DC) ATAC-seq peaks overlap a direct alignment between zebrafish and mouse. Indirectly conserved (IC) ATAC-seq peaks do not overlap a direct alignment, but are projected with a score > 0.99, i.e. either overlapping or very close to a multi-species anchor. The remaining peaks are classified as non-conserved (NC). A score of 0.99 means that the sum of the distances from peak to anchor points is < 150 bps considering all intermediate species in the optimal species path.

### Motif search

Motif hits were computed using seqPattern (Haberle et al., 2021, doi: https://doi.org/10.18129/B9.bioc.seqPattern) on a set of position weight matrices for 120 transcription factor families. Regions with scores of > 85 were considered as motif hits.

## Acknowledgements

We thank Max Haussler at UCSC, and Daniel Zerbino at EBI for facilitating access of DANIO-CODE track hubs in the UCSC and Ensembl genome browsers respectively. We thank ZFIN for hosting the DANIO-CODE DCC and raw data. We thank Julia Horsefield for creating the DANIO-CODE logo. We thank data producers (for list of laboratories visit the DANIO-CODE DCC) who directly uploaded data and provided metadata directly. We thank DNANexus for providing compute time for the reprocessing of public datasets.

We thank our main funders, the Horizon 2020 MSCA-ITN project ZENCODE-ITN by the European Commission to FM, BL, CD, JMV and BP (Grant agreement no: 643062). FM, BL, and FW thank BBSRC support (DanioPeaks, P61715). FM and BL thank support by the Wellcome Trust (Joint-Investigator award 106955/Z/15/Z). We thank SNP&SEQ Technology Platform in Uppsala, Sweden (CAGE sequencing) and DB was awarded the Rutherford Fund Fellowship.

## Supplementary Figures

**Supplementary Figure 1.**
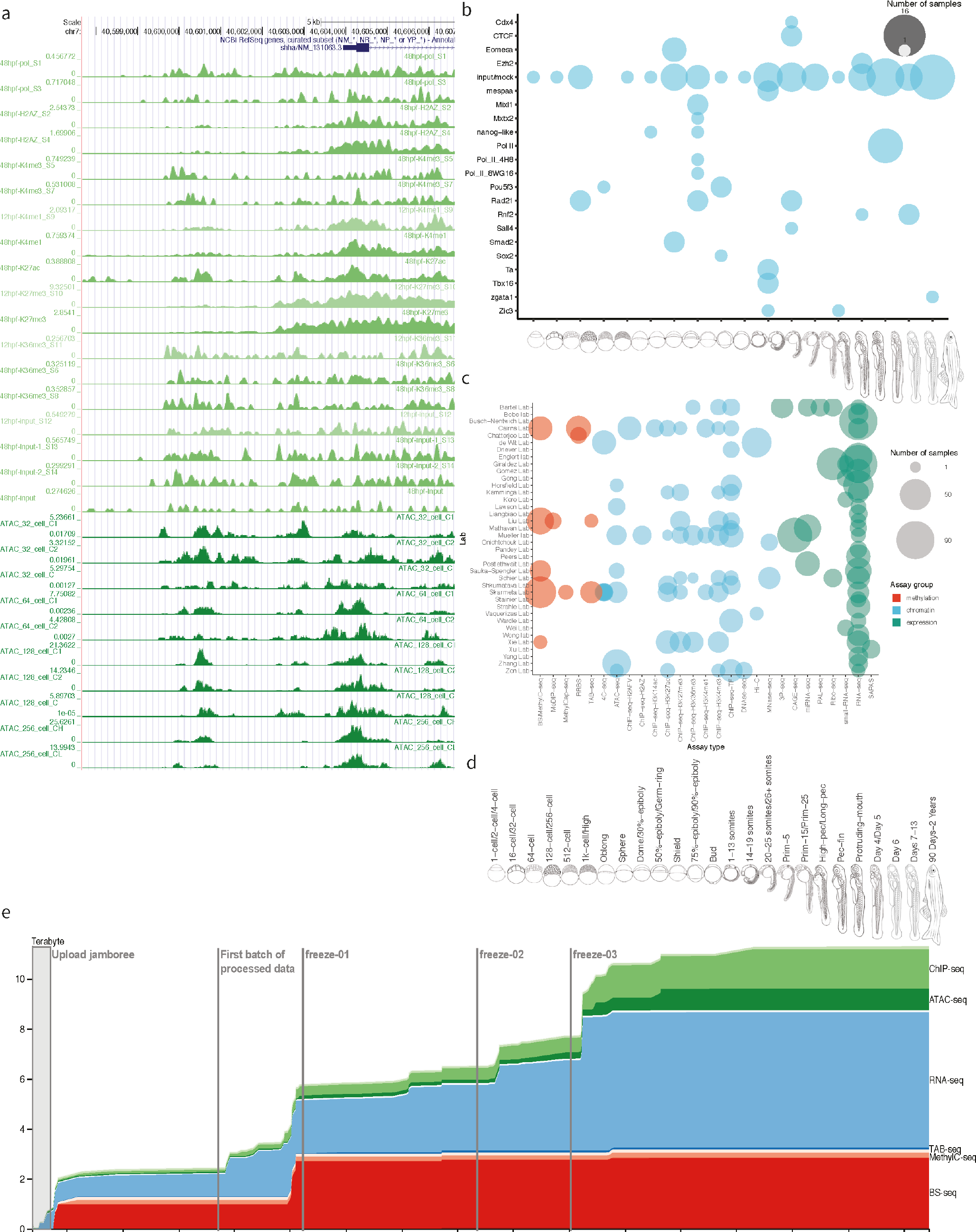
**a**, Tracks of representative examples of unpublished datasets in a UCSC Genome Browser session including CAGE, ATAC, and ChIP datasets generated by DANIO-CODE laboratories. Promoter region of developmental regulator *shha* gene is shown. **b**, DCC Data availability summary for ChIP with antibodies against Pol II, CTCF and transcription factors as indicated. Stages and stage ranges are indicated on the X axis, the transcription factor occupancy detected is listed on the Y axis. **c**, Data producers and data types matrix indicating the data producer lab (Y axis) and the type of data (X axis). **d**, Data acquisition evolution in the DCC.

**Supplementary Figure 2.**
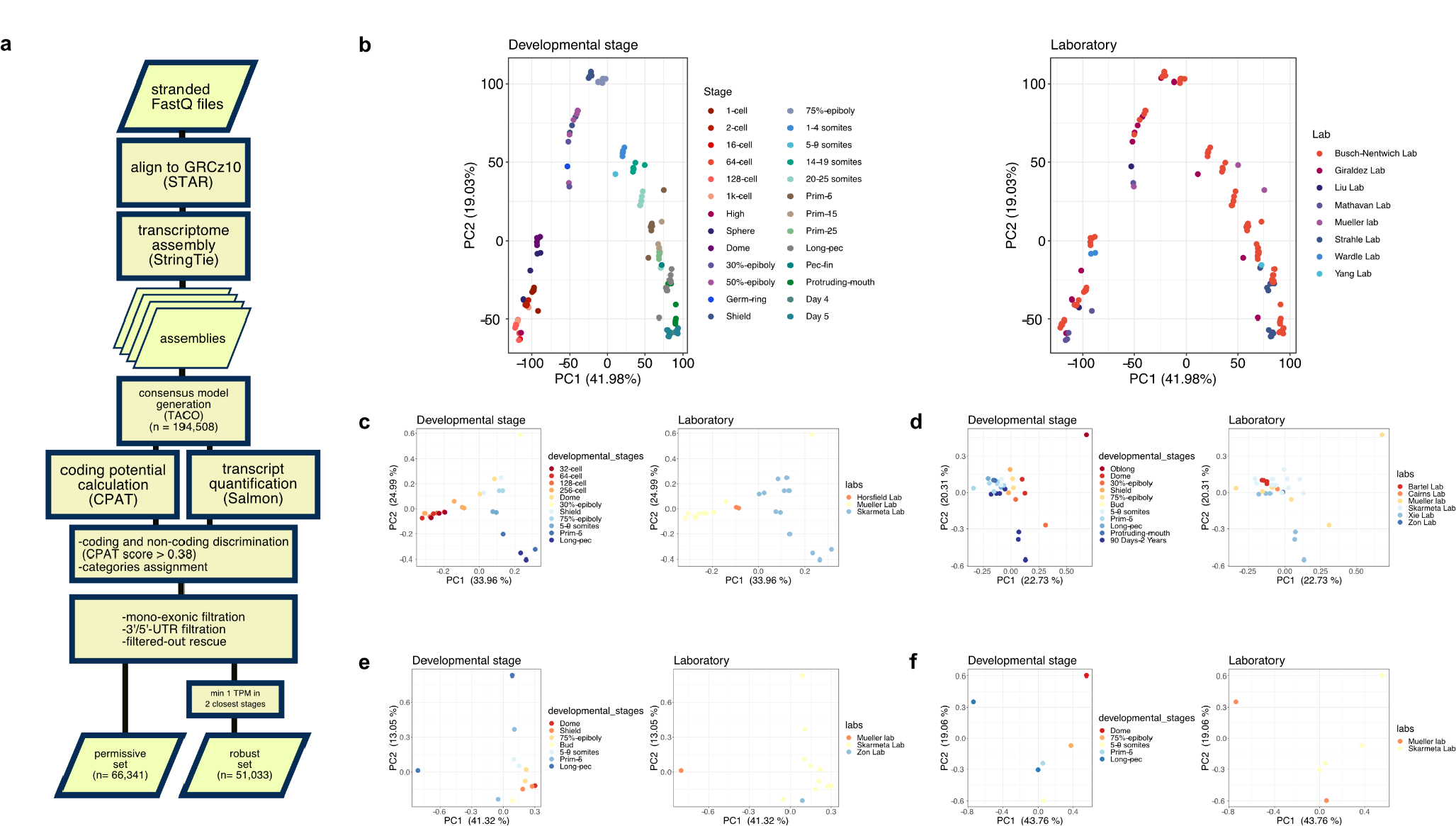
**a**, RNA-seq data processing pipeline, **b**, PCA of CAGE-supported RNA seq (n=14,885), labeled by stages (left) and labs (right). PCA of consensus open regions (cPADREs; see Suppl. Figure 6a) of **c**, ATAC-seq **d**, H3K4me **e**, H3K27ac **f**, and H3K4me1 (h). For H3K4me3 regions were restricted to the promoter-overlapping set.

**Supplementary Figure 3.**
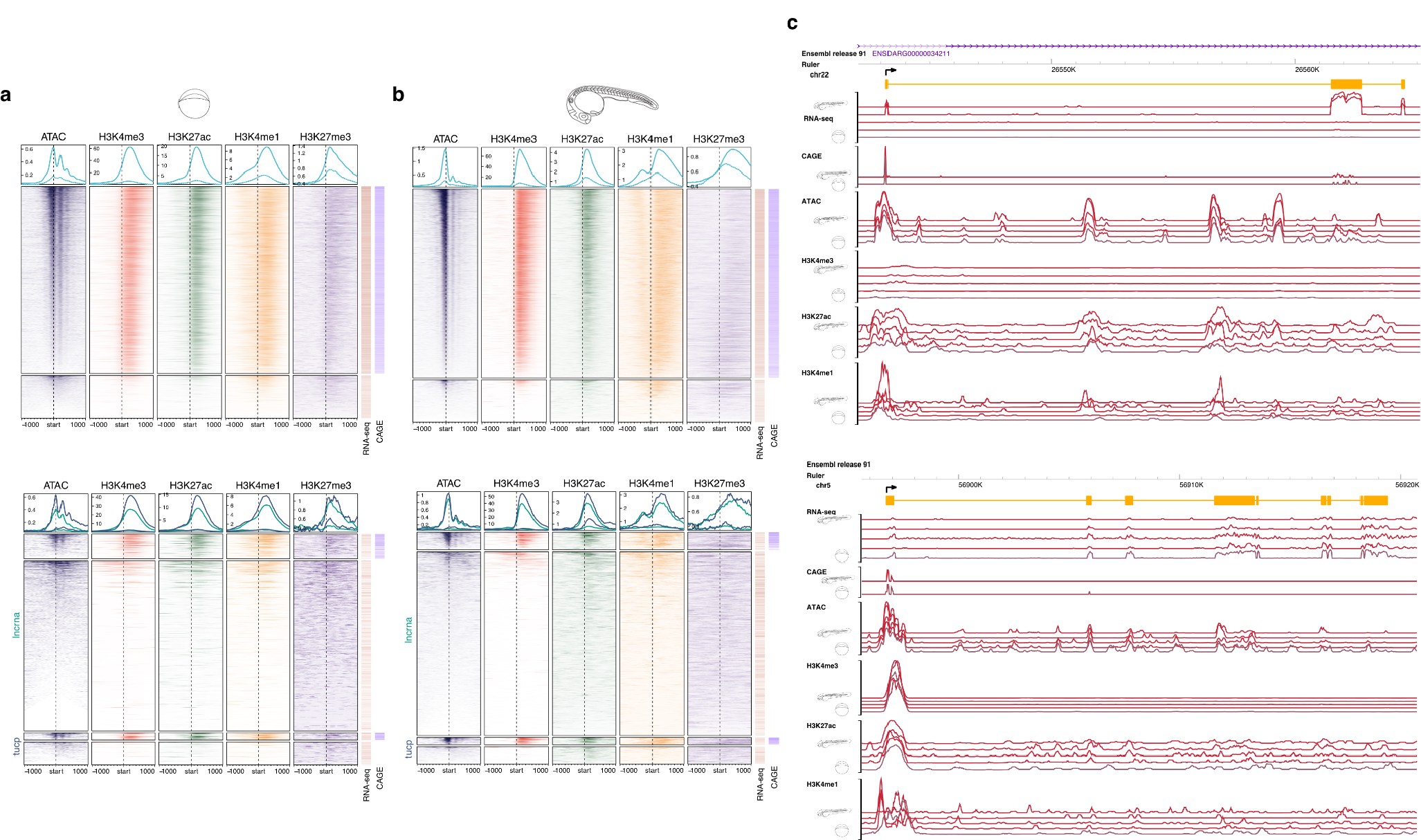
**a, b**, Aggregation plots and heatmaps of open chromatin and epigenetic features of annotated transcripts and CAGE-seq validation of TSSs (bars on the right of each panel) are shown for the Dome (left) and Prim-5 (right) stages. Top panels show protein coding genes (n=14,246, of which 11,586 are supported by CAGE for the Dome stage; n=16180, of which 13,200 are supported by CAGE for the Prim-5 stage) and bottom panels show lncRNA (n=2,308, of which 296 are supported by CAGE for the Dome stage; n=2,774, of which 246 are supported by CAGE for prim-5) and TUCP (n=351, of which 81 are supported by CAGE for the Dome stage; n=331, of which 101 are supported by CAGE for the Prim-5 stage) genes **c,** Example screenshot of novel lncRNA(top), and novel TUCP (bottom) transcripts and epigenomic features.

**Supplementary Figure 4.**
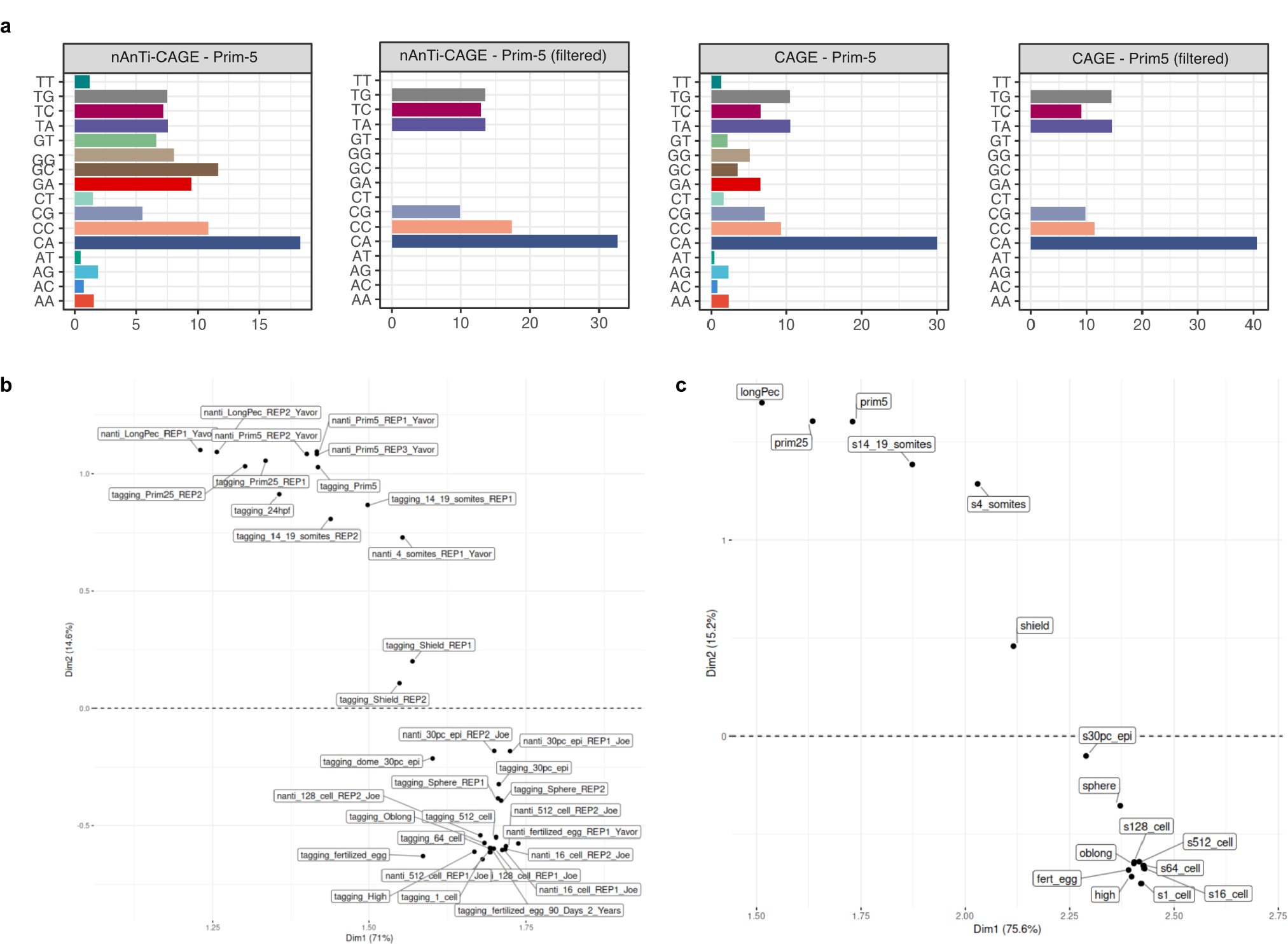
**a**, dinucleotide frequency analysis of TSS clusters before and after canonical filtering, **b**, PCA correlation analysis of TSS activity detected by two CAGE protocols, **c**, PCA of 27,781 consensus promoters.

**Supplementary Figure 5.**
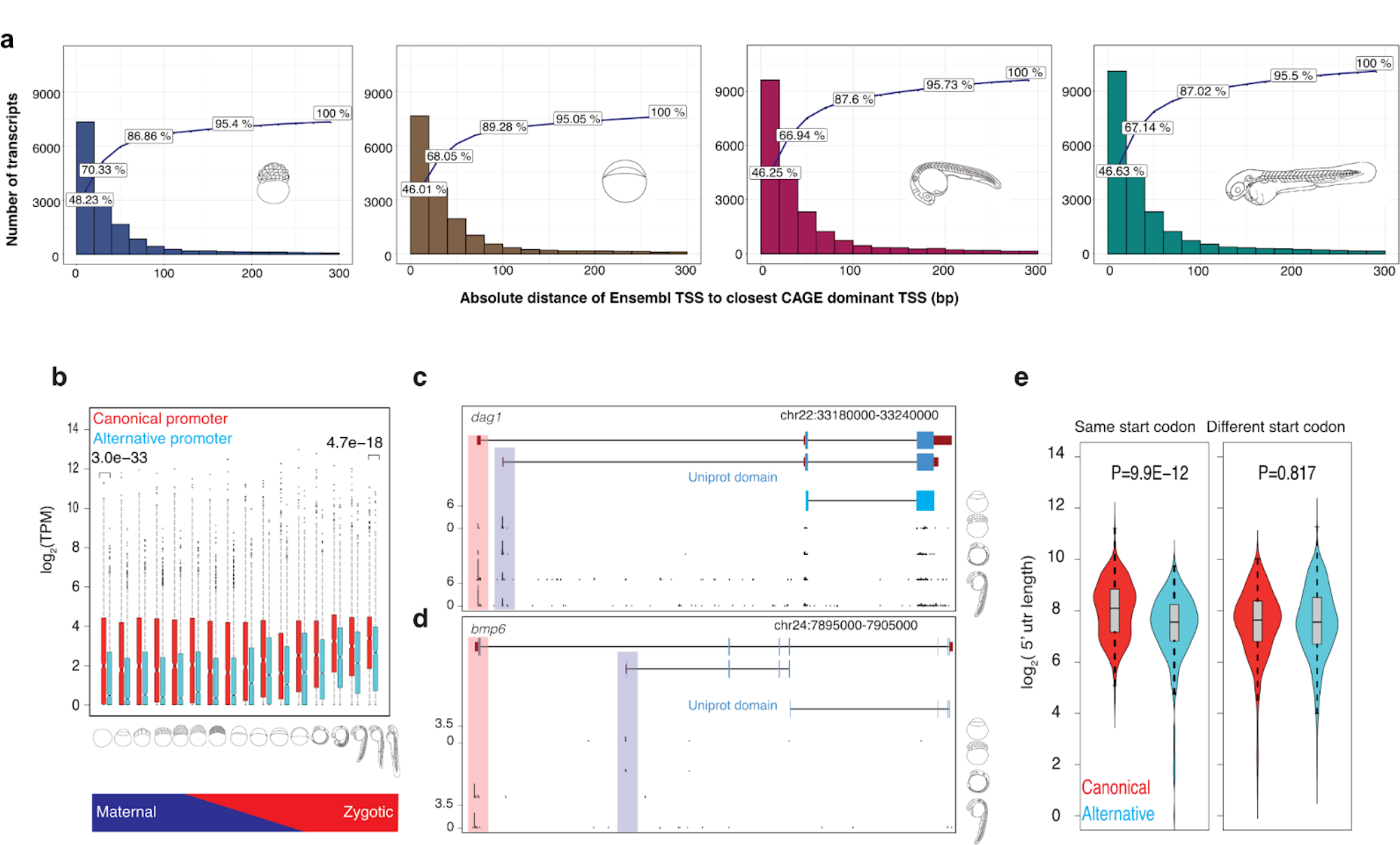
**a**, Frequency distribution of ensembl transcripts 5’ ends binned according to distance (bp) from CAGE dominant peak as indicated on X axis. Cumulative frequency depicted by line. Developmental stages are indicated by embryo symbols. **b,** Box plot shows the expression levels of canonical and alternative promoters across 16 developmental stages. P-values denote the significant difference in expression levels between canonical and alternative promoters during two stages at fertilized-egg (P=3.0E-33; t-test two-sided) and long pec (P=4.7E-18; t-test two-sided). **c,** A UCSC browser screenshot of the gene dag1 shows the alternative promoter (highlighted in cyan) is upstream of the start codon (pointed by arrow), thus altering only 5’UTR but not protein. The numbers on the y-axis represent the normalized tags per million (TPM) of CAGE tags. The Uniprot domain track denotes the annotated protein domain in the Uniprot database. **d,** A UCSC browser screenshot of the gene *bmp6* shows the alternative promoter (highlighted in cyan) is downstream of the start codon (pointed by arrow) and alters the N terminal of the protein. **e,** Bean plots show 5’ UTR length of transcripts supported by canonical and alternative promoters which are grouped based on the usage of the same or different start codon. Alternative promoters that utilized the same start codon have significantly shorter 5’ UTR length (t-test, two-sided).

**Supplementary Figure 6.**
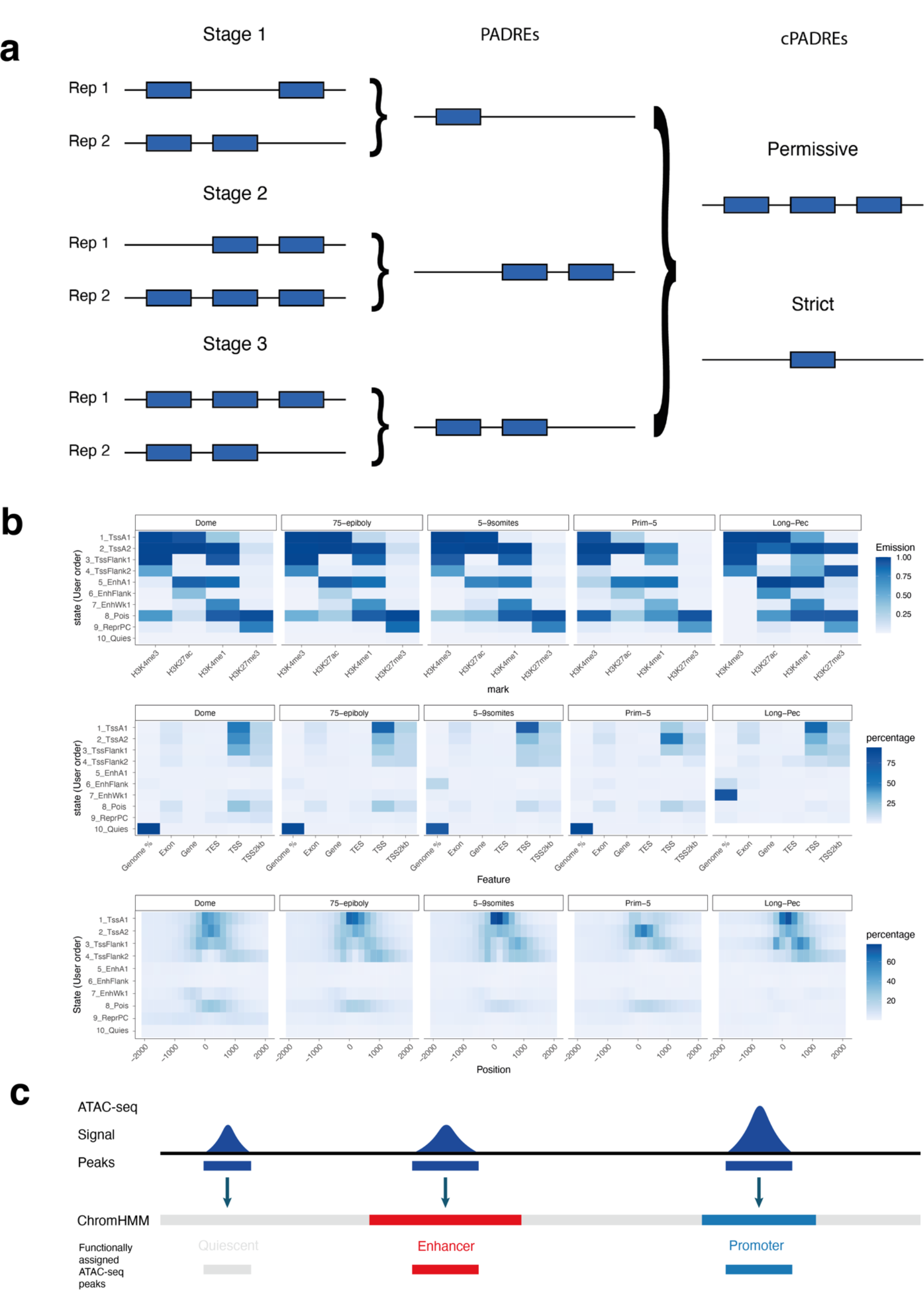
**a**, Schematic representation of PADREs and consensus PADREs (cPADREs) definition. PADREs are defined as ATAC-seq peaks reproducible between replicates. We defined cPADREs as a union of PADREs (permissive) and further filtered them with a requirement that a region needs to be open in at least 2 neighbouring developmental stages. The later set was termed a strict cPADRE set, and used for the analysis in this paper **b,** Top: Occurrence probabilities of chromatin marks in each obtained ChromHMM state for five developmental stages. Middle: Enrichment of genomic features in each state for five developmental stages. Bottom: Occurrence of each state +/- 2kb around RefSeq TSS for five developmental stages. The states function was manually assigned. 1_TssA1, 2_TssA2 = Active TSS, 3_TssFlank1, 4_TssFlank2 = TSS Flanking region, 5_EnhA1 = Active enhancer, 6_EnhFlank = Weak enhancer, 7_EnhWk1 = Primed enhancer, 8_Pois = Poised elements, 9_PcRep = Polycomb repressed regions, 10_Quies = Quiescent state. **c,** Functional assignment of PADREs using ChromHMM genome segmentation

**Supplementary Figure 7.**
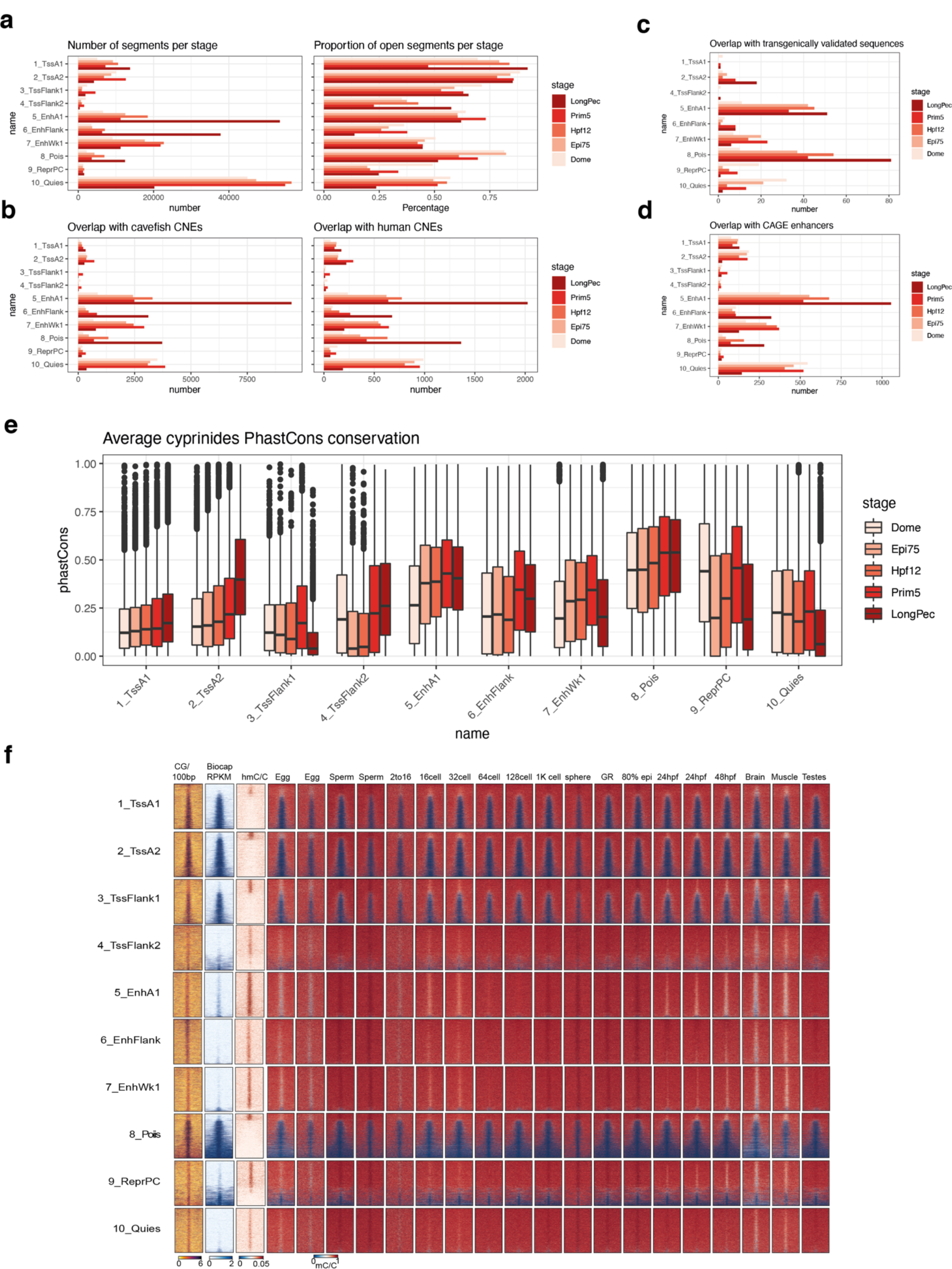
**a**, Left: Number of PADREs assigned to each chromatin state for every developmental stage. Right: Proportion of ChromHMM states present in PADREs for each stage. **b,** Number of annotated PADREs overlapping Mexican cavefish (left) and human (right) CNEs for each stage. **c,** Number of annotated PADREs overlapping transgenically validated enhancers for each stage. **d,** Number of annotated PADREs overlapping CAGE-defined eRNAs for each stage. **e,** phastCons scores distribution of annotated PADREs for each stage. **f,** Methylation profile throughout development of annotated PADREs at the Prim-5 stage

**Supplementary Figure 8.**
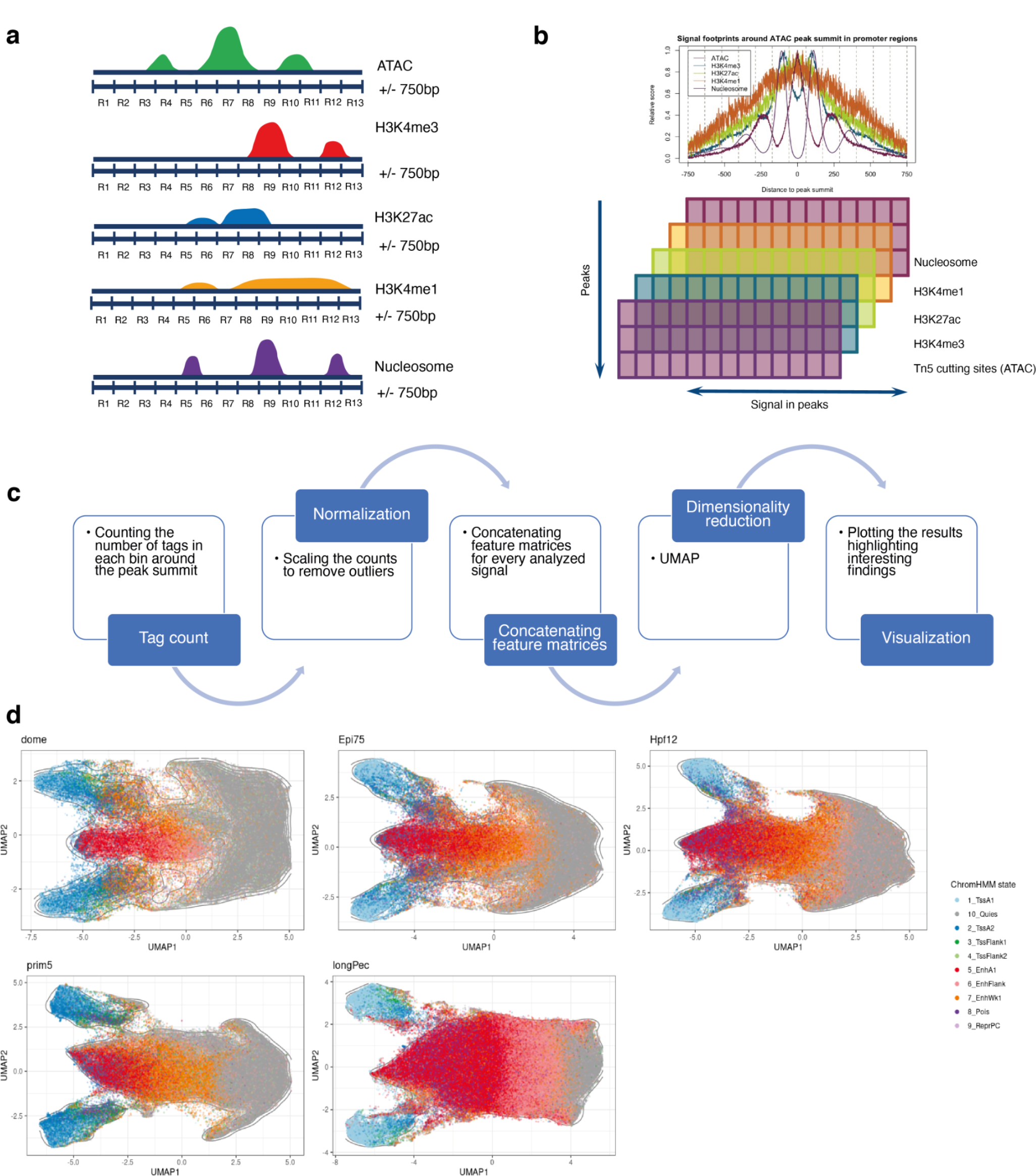
**a-c**, Schematic representation of UMAP visualization of PADREs (for details, see Methods). R1-13 represent bins used to make the model. **d,** UMAP plot of annotated PADREs for each developmental stage analysed

**Supplementary Figure 9.**
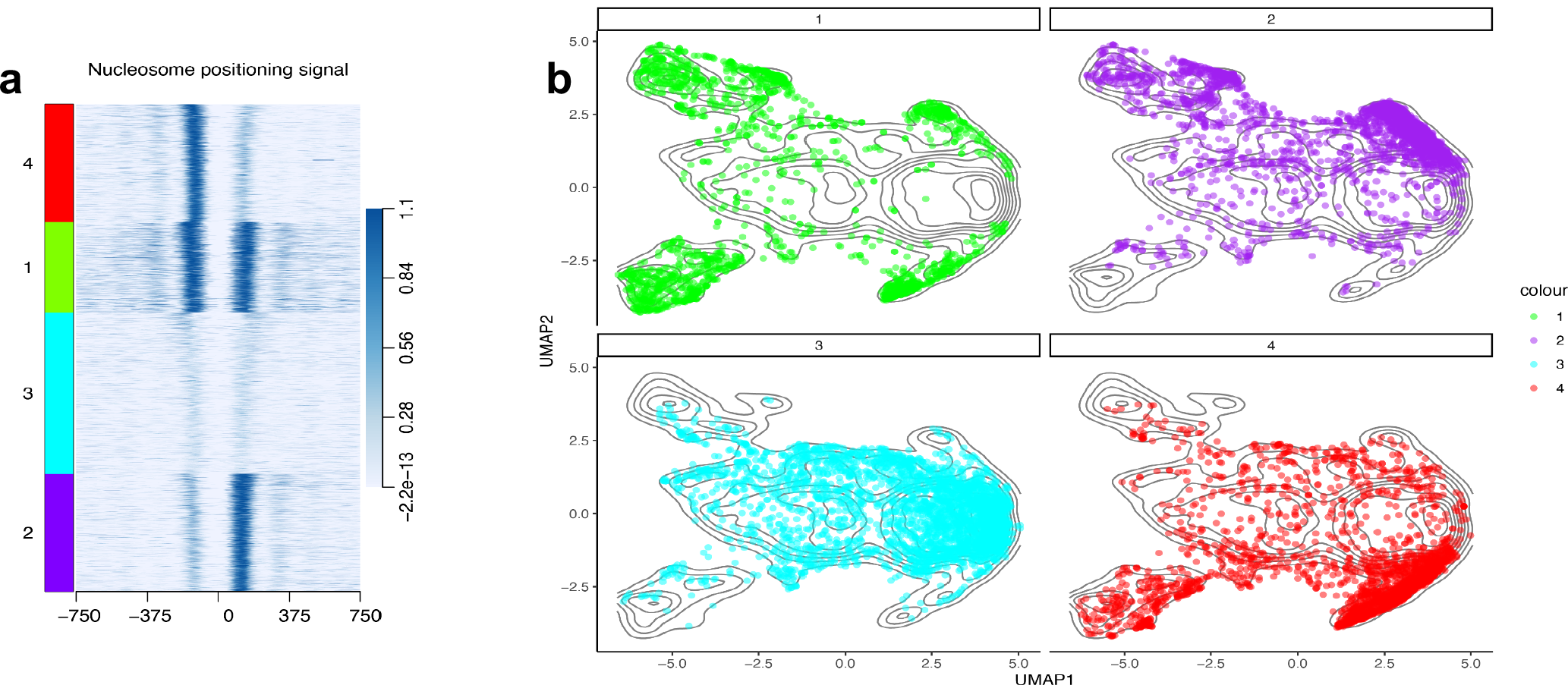
Nucleosome position around PADREs containing the CTCF motif. **a,** Heatmap with nucleosome position signal PADREs containing the CTCF motif, divided into four groups using the k-means algorithm. **b,** UMAP position of PADREs belonging to groups from **a**.

**Supplementary figure 10.**
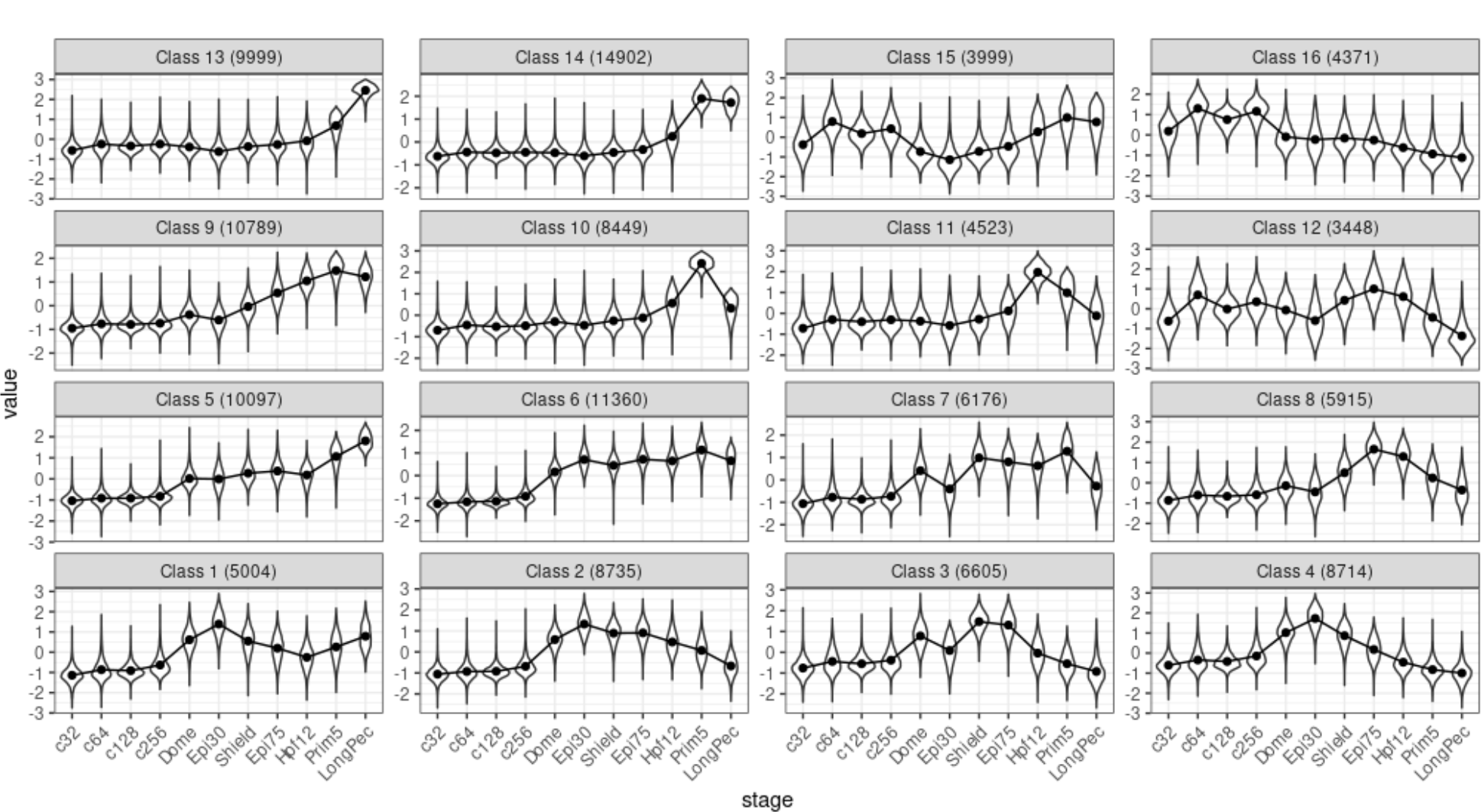
Openness of distal cPADREs throughout development in all SOM classes at stages as indicated. Numbers in brackets indicate the number of elements.

**Supplementary Figure 11.**
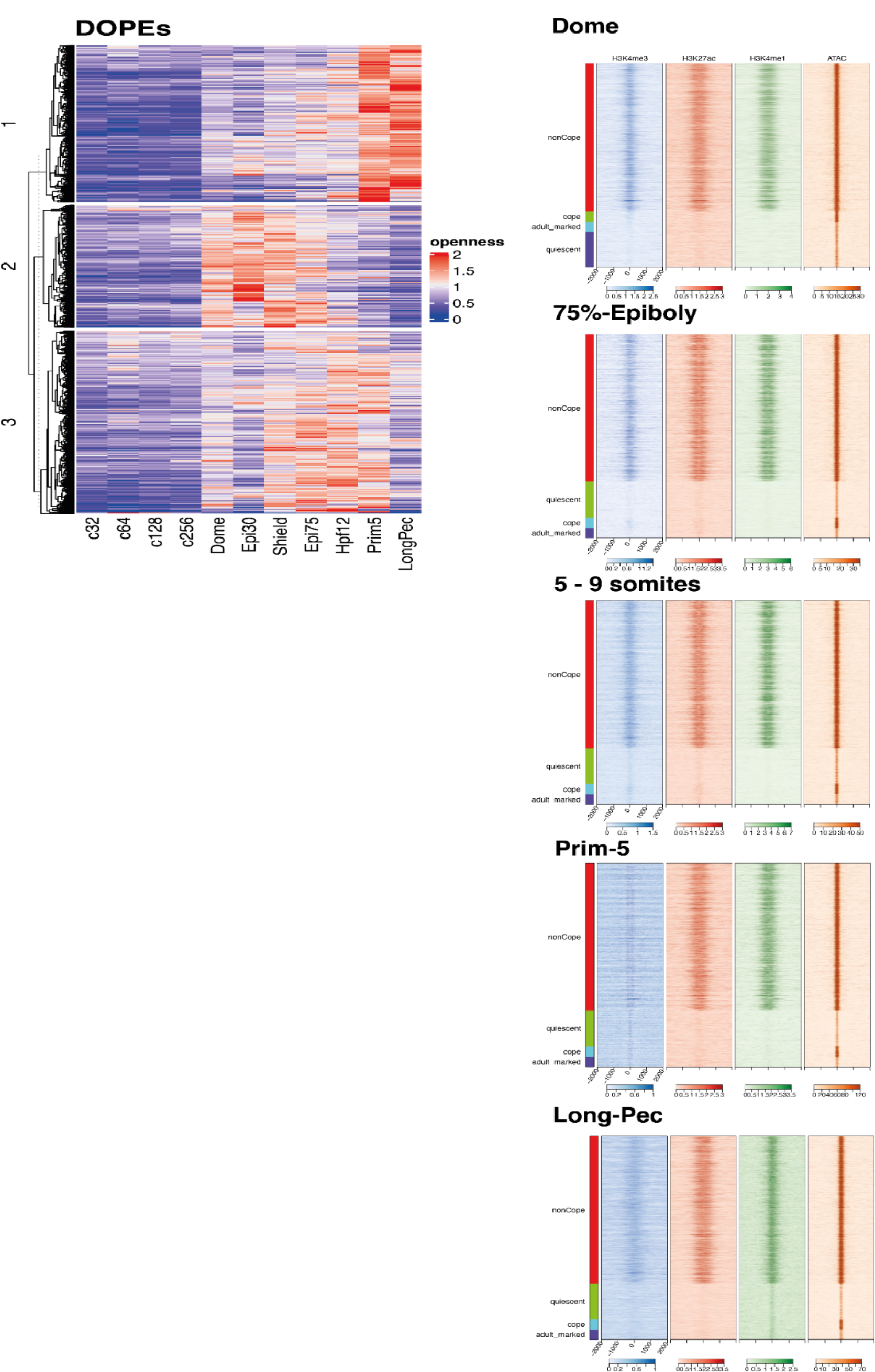
**a**, DOPEs and their chromatin openness throughout development. DOPEs were clustered into three groups based on the time of chromatin opening using *k*-means clustering. **b,** Signal heatmaps of H3K4me3, H3K27ac, H3K4me1, and ATAC for COPEs, DOPEs, DOPEs active in adult tissues, and other constitutive elements throughout development aligned to the centre of open chromatin.

**Supplementary Figure 12.**
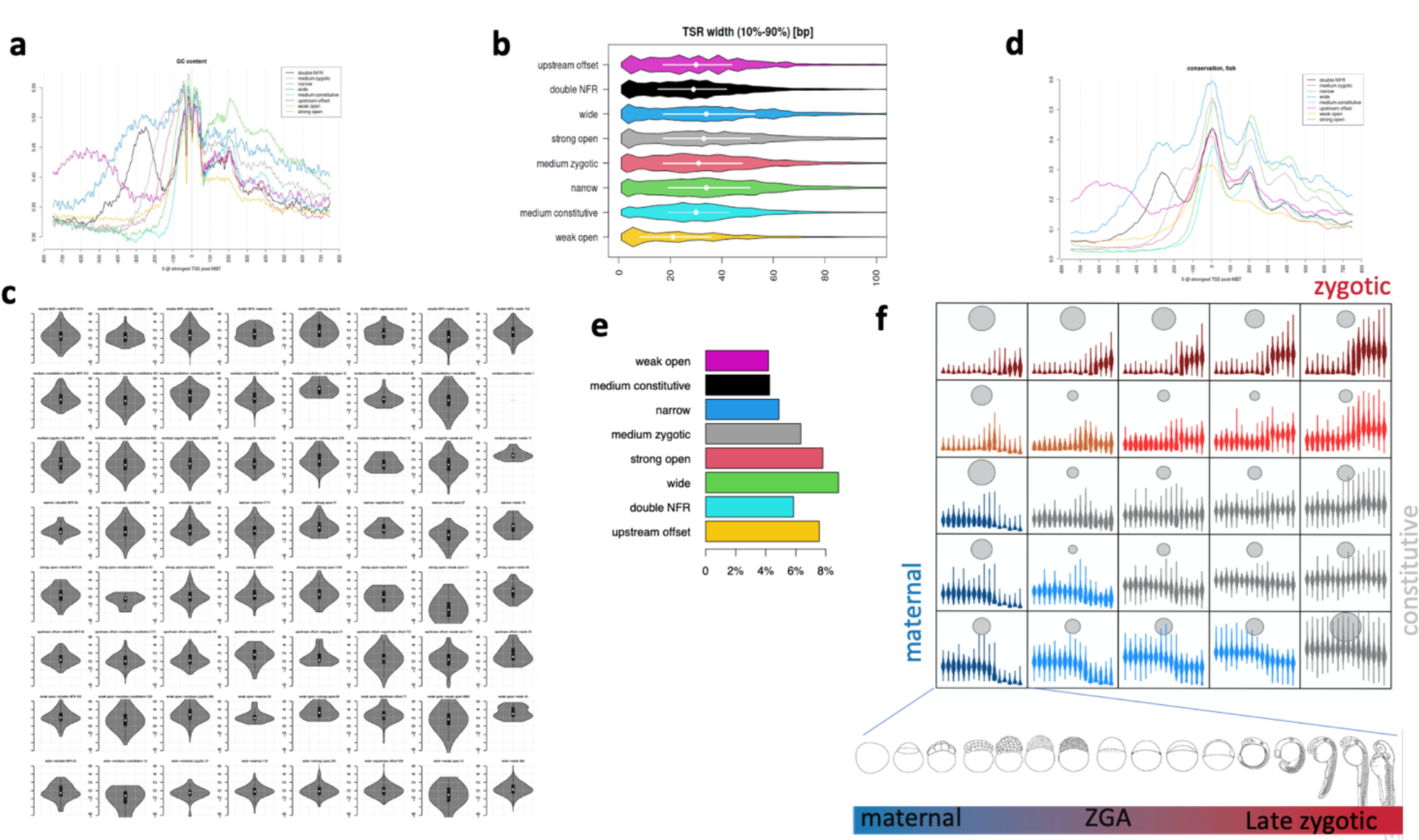
**a**, Promoters of each class are enriched in GC content. GC content stays high until +50bp when it abruptly switches into the WW oscillating region (nucleosome positioning signal). The fine structure is similar in each class. The nucleosome positioning oscillation is diminished around the nucleosome dyad (+120bp), and then it reappears in the second half of the nucleosome-wrapped fragment. **b**, Transcription start region (TSR) width in base pairs measured by CAGE signal distribution, incorporating the central 80% of CAGE expression. **c**, Expression dynamics of promoters transitioning between classes at the Dome and Prim-5 stages (violin plots of log fold changes). **d,** Aggregated teleost sequence conservation plotted against the dominant transcription start site (0). Sequence conservation always peaks exactly at the TSS. There are additional conservation peaks visible downstream between nucleosome linker regions. In three cases where there are open regions upstream, conservation follows openness upstream. Generally, conservation follows openness, suggesting purifying selection acting on regulatory sites. **e,** Bar chart of % proportion of promoter classes containing target genes of Gene regulatory Block (GRBs, see text associated with Figure 5). **f,** Classification of promoter expression during development with self-organising maps (SOM). 5×5 diagrams contain violin plots with stage-by-stage expression levels. Blue to red spectrum indicates maternal to zygotic expression dynamics of promoter clusters. Gray circles indicate the number of promoters per cluster. Stages of development are symbolised below the SOM array.

**Supplementary Figure 13.**
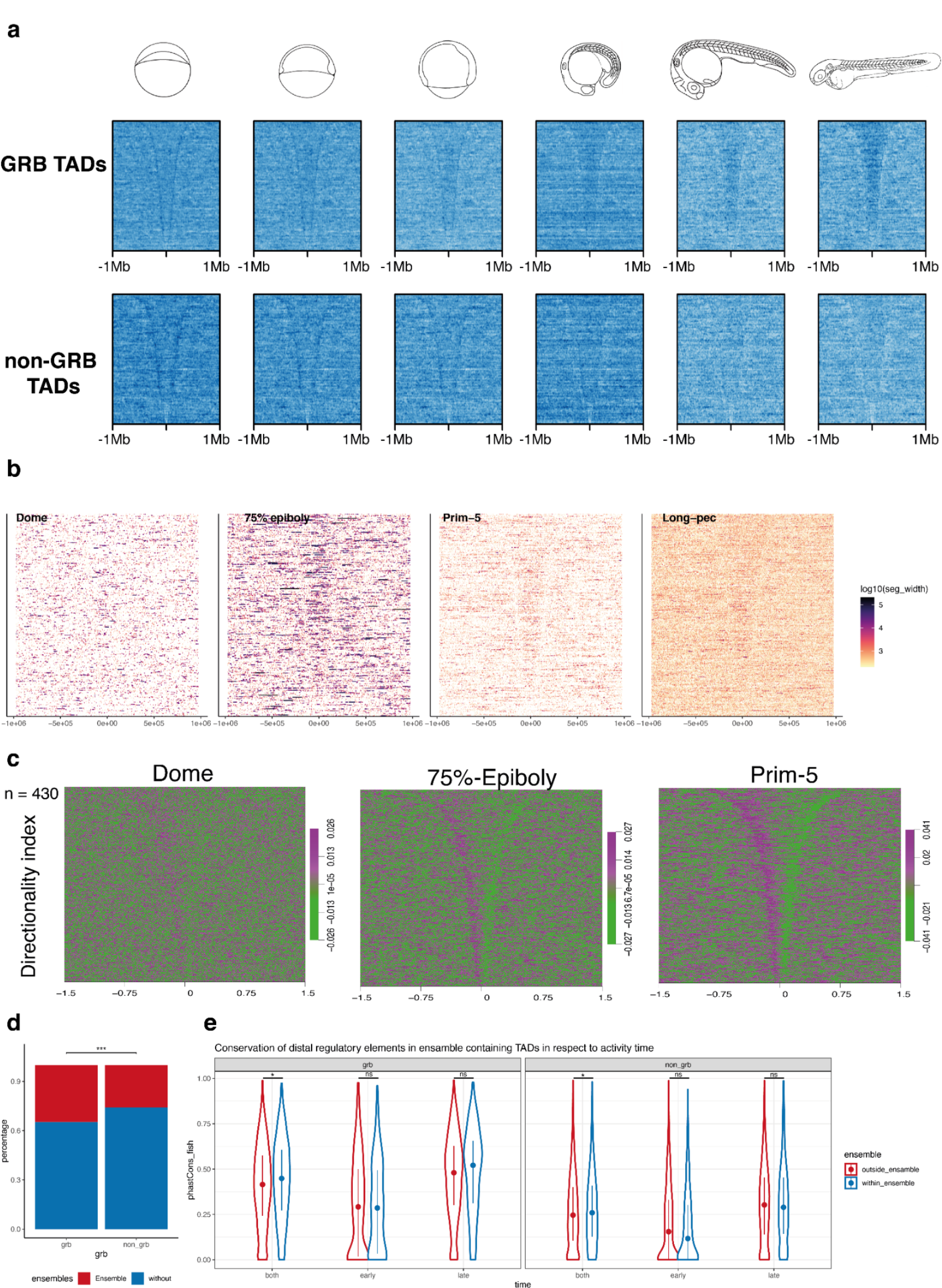
**a**, ATAC-seq signals in GRB (top) and non-GRB (bottom) TADs throughout development. TADs are ordered in a descending order from the top of the heatmap.**b,** Enhancer-associated ChromHMM segments in GRB TADs throughout development. TADs are ordered in a descending order from the top of the heatmap. Segments are coloured based on the logarithm of their length. Early stages are dominated by fewer large blocks, which start to be enriched within TADs only at 75%-Epiboly. In late stages, short segments are distributed uniformly throughout the entire TAD length. **c,** Directionality index in GRB TADs throughout development. **e,** phastCons scores of distal PADREs within or outside TADs. PADREs are separated on the x-axis based on the stage they are open, and plots are separated based on type of TAD.

**Supplementary Figure 14.**
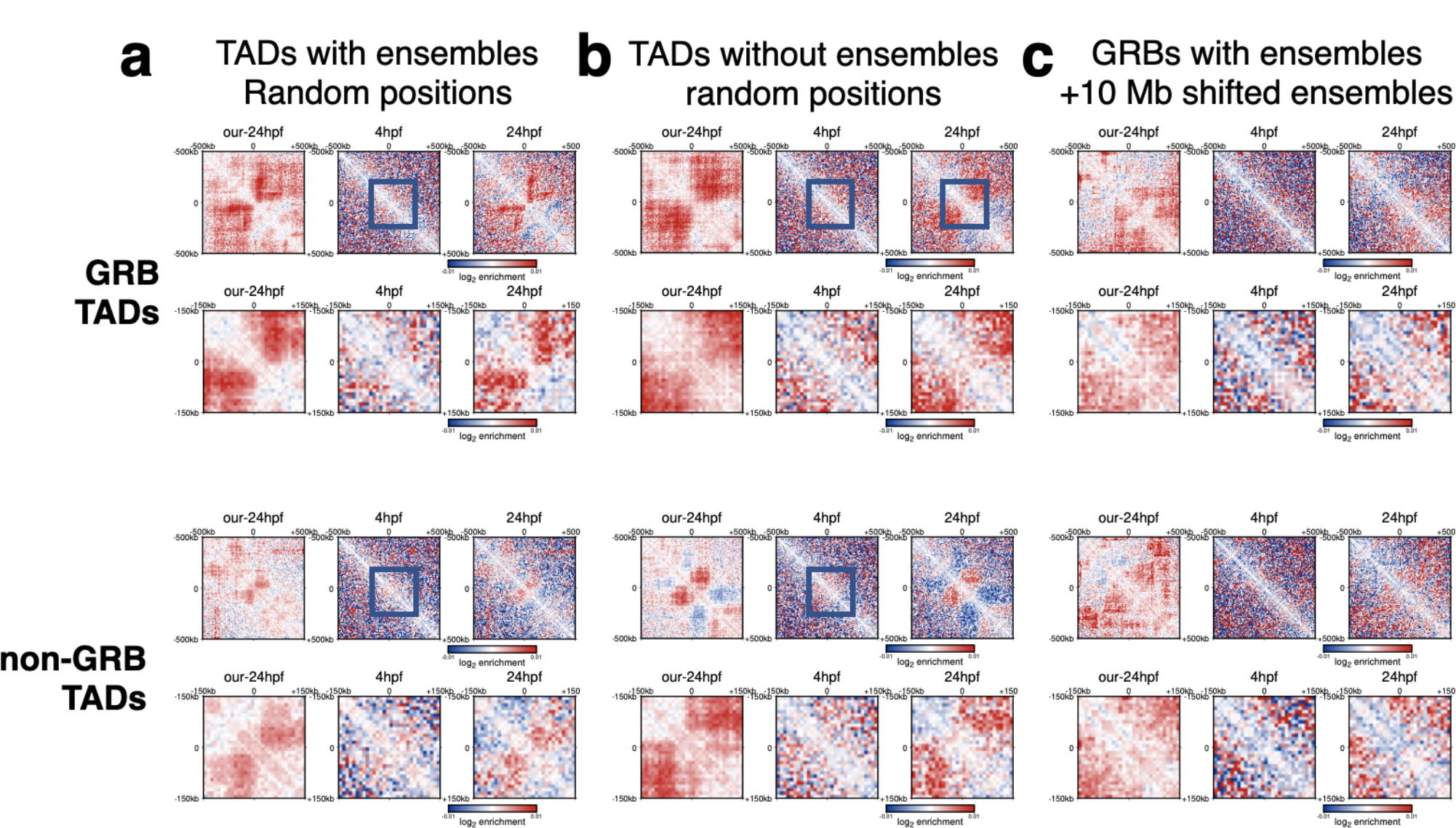
**a-c**, Controls for contact enrichment around H3K27ac ensembles. All regions were downsampled to n=56 to match the number of 50 kb - 150 kb size ensembles. Labels are as in Figure 5g. The controls included random positions within the same TAD (**a**), random positions within TADs without ensembles (**b**), and 10MB shifted positions (**c**), for GRB TADs (top row) and non-GRB TADs (bottom row). The controls include published data for the Prim-5 stage, as well as new, unpublished data with higher resolution (Prim 5)..

**Supplementary Figure 15.**
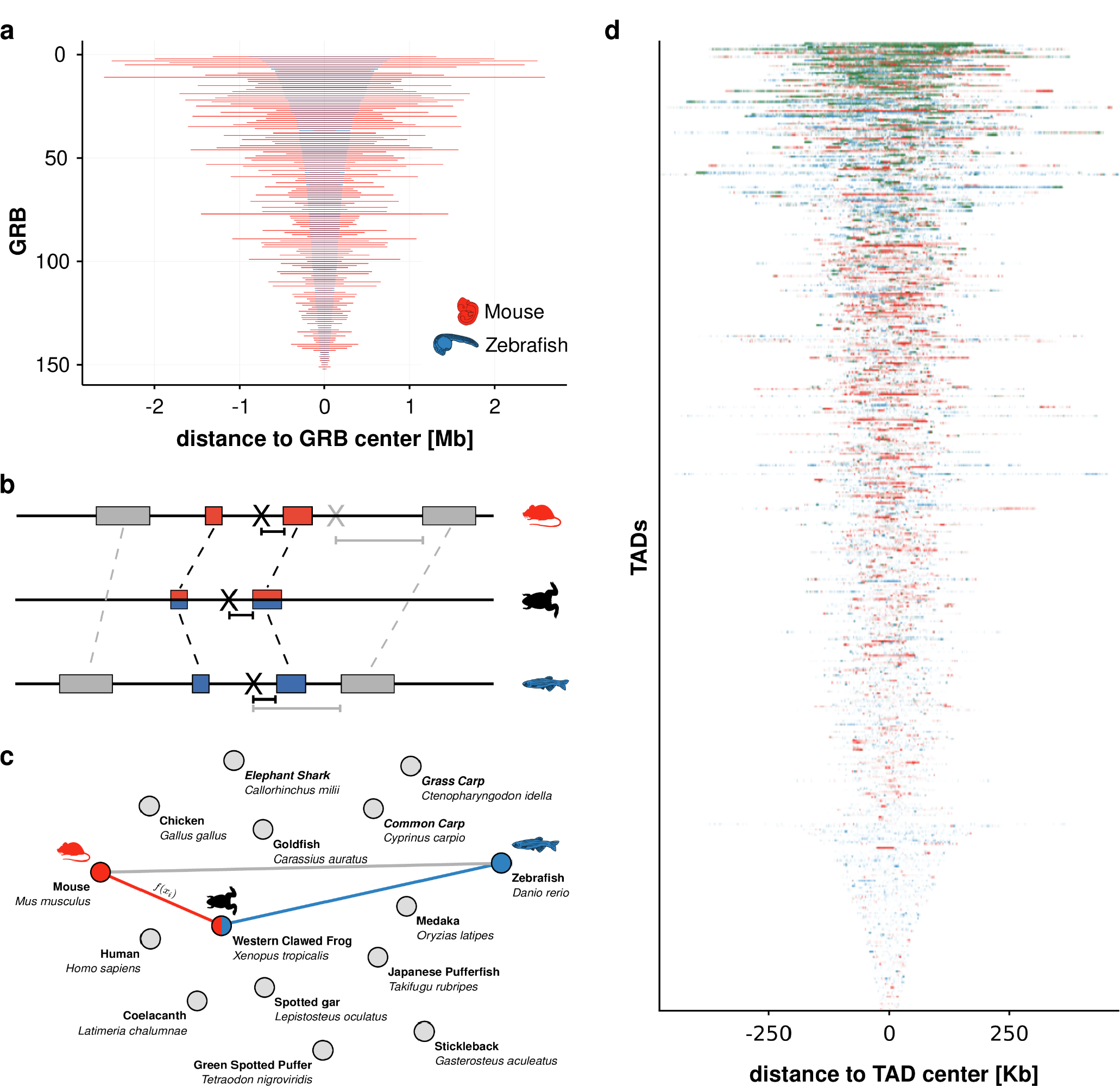
**a**, Schematic illustration of the projection of an example genomic location X between zebrafish and mouse by interpolation using the direct alignments (grey rectangles) and the alignments via a bridging species (blue and red rectangles, Xenopus in this example). projections are indicated as a black X in the respective species). Dashed lines connect pairwise sequence alignments. The projected locations of X in mouse are indicated in grey (direct alignments) and black (via bridging species). **b,** Example graph comprising 15 species (nodes). For any genomic location, the shortest path through the species graph yields the combination of species which maximizes projection accuracy. **c,** Comparison of sizes of genomic sequences covering orthologous GRB-containing TADs. TADs are ranked by size, largest on top. **d,** H3K27me3 overlap profiles of all GRB TADs. TADs are ordered by their relative amount of shared signal. Bins are in the original genomic order.

## References

1. Adamson, K.I., Sheridan, E. & Grierson, A.J. Use of zebrafish models to investigate rare human disease. Journal of Medical Genetics 55, 641–649 (2018).

2. Bradford, Y.M. et al. Zebrafish Models of Human Disease: Gaining Insight into Human Disease at ZFIN. ILAR Journal 58, 4–16 (2017).

3. Patton, E.E. & Tobin, D.M. Spotlight on zebrafish: the next wave of translational research.Disease Models & Mechanisms 12(2019).

4. Teame, T. et al. The use of zebrafish (Danio rerio) as biomedical models. Animal Frontiers 9, 68–77 (2019).

5. Howe, D.G. et al. The Zebrafish Model Organism Database: new support for human disease models, mutation details, gene expression phenotypes and searching. Nucleic Acids Research 45, D758–D768 (2017).

6. Ablain, J. & Zon, L.I. Of fish and men: using zebrafish to fight human diseases. Trends in Cell Biology 23, 584–586 (2013).

7. Howe, K. et al. The zebrafish reference genome sequence and its relationship to the human genome. Nature 496, 498–503 (2013).

8. Kettleborough, R.N.W. et al. A systematic genome-wide analysis of zebrafish protein-coding gene function. Nature 496, 494–497 (2013).

9. Bogdanovic, O. et al. Dynamics of enhancer chromatin signatures mark the transition from pluripotency to cell specification during embryogenesis. Genome Res 22, 2043–53 (2012).

10. Vastenhouw, N.L. et al. Chromatin signature of embryonic pluripotency is established during genome activation. Nature 464, 922–926 (2010).

11. Murphy, P.J., Wu, S.F., James, C.R., Wike, C.L. & Cairns, B.R. Placeholder Nucleosomes Underlie Germline-to-Embryo DNA Methylation Reprogramming. Cell 172, 993–1006.e13 (2018).

12. Haberle, V. et al. Two independent transcription initiation codes overlap on vertebrate core promoters. Nature 507, 381–385 (2014).

13. Nepal, C. et al. Dual-initiation promoters with intertwined canonical and TCT/TOP transcription start sites diversify transcript processing. Nat Commun 11, 168 (2020).

14. Zhao, L., Wang, L., Chi, C., Lan, W. & Su, Y. The emerging roles of phosphatases in Hedgehog pathway. Cell Communication and Signaling 15(2017).

15. Bazzini, A.A., Lee, M.T. & Giraldez, A.J. Ribosome Profiling Shows That miR-430 Reduces Translation Before Causing mRNA Decay in Zebrafish. Science 336, 233–237 (2012).

16. Lee, M.T. et al. Nanog, Pou5f1 and SoxB1 activate zygotic gene expression during the maternal-to-zygotic transition. Nature 503, 360–364 (2013).

17. Leichsenring, M., Maes, J., Mössner, R., Driever, W. & Onichtchouk, D. Pou5f1 Transcription Factor Controls Zygotic Gene Activation In Vertebrates. Science 341, 1005–1009 (2013).

18. Bogdanović, O. et al. Active DNA demethylation at enhancers during the vertebrate phylotypic period. Nature Genetics 48, 417–426 (2016).

19. Potok, M.E., Nix, D.A., Parnell, T.J. & Cairns, B.R. Reprogramming the Maternal Zebrafish Genome after Fertilization to Match the Paternal Methylation Pattern. Cell 153, 759–772 (2013).

20. Jiang, L. et al. Sperm, but Not Oocyte, DNA Methylome Is Inherited by Zebrafish Early Embryos. Cell 153, 773–784 (2013).

21. Satija, R., Farrell, J.A., Gennert, D., Schier, A.F. & Regev, A. Spatial reconstruction of single-cell gene expression data. Nature Biotechnology 33, 495–502 (2015).

22. Kikuta, H. et al. Genomic regulatory blocks encompass multiple neighboring genes and maintain conserved synteny in vertebrates. Genome Research 17, 545–555 (2007).

23. Gehrig, J. et al. Automated high-throughput mapping of promoter-enhancer interactions in zebrafish embryos. Nat Methods 6, 911–6 (2009).

24. Rada-Iglesias, A. et al. A unique chromatin signature uncovers early developmental enhancers in humans. Nature 470, 279–283 (2010).

25. Spieler, D. et al. Restless Legs Syndrome-associated intronic common variant in Meis1 alters enhancer function in the developing telencephalon. Genome Research 24, 592–603 (2014).

26. Encode Project Consortium. An integrated encyclopedia of DNA elements in the human genome. Nature 489, 57–74 (2012).

27. Dixon, J.R. et al. Chromatin architecture reorganization during stem cell differentiation. Nature 518, 331–336 (2015).

28. Kundaje, A. et al. Integrative analysis of 111 reference human epigenomes. Nature 518, 317–330 (2015).

29. Gerstein, M.B. et al. Integrative Analysis of the Caenorhabditis elegans Genome by the modENCODE Project. Science 330, 1775–1787 (2010).

30. Roy, S. et al. Identification of Functional Elements and Regulatory Circuits by Drosophila modENCODE. Science 330, 1787–1797 (2010).

31. Yang, H. et al. A map of cis-regulatory elements and 3D genome structures in zebrafish. Nature 588, 337–343 (2020).

32. Tan, H., Onichtchouk, D. & Winata, C. DANIO-CODE: Toward an Encyclopedia of DNA Elements in Zebrafish. Zebrafish 13, 54–60 (2016).

33. Hortenhuber, M., Mukarram, A.K., Stoiber, M.H., Brown, J.B. & Daub, C.O. *-DCC: A platform to collect, annotate, and explore a large variety of sequencing experiments. Gigascience 9(2020).

34. Encode Project Consortium et al. Expanded encyclopaedias of DNA elements in the human and mouse genomes. Nature 583, 699–710 (2020).

35. Fantom Consortium the Riken Pmi Clst et al. A promoter-level mammalian expression atlas. Nature 507, 462–70 (2014).

36. Kent, W.J. et al. The Human Genome Browser at UCSC. Genome Research 12, 996–1006 (2002).

37. Donlin, M.J. Using the Generic Genome Browser (GBrowse). Curr Protoc Bioinformatics Chapter 9, Unit 9 9 (2009).

38. Howe, K.L. et al. Ensembl 2021. Nucleic Acids Res 49, D884–D891 (2021).

39. Li, D., Hsu, S., Purushotham, D., Sears, R.L. & Wang, T. WashU Epigenome Browser update 2019. Nucleic Acids Res 47, W158–W165 (2019).

40. McGarvey, A.C. et al. (2020).

41. Pauli, A. et al. Systematic identification of long noncoding RNAs expressed during zebrafish embryogenesis. Genome Res 22, 577–91 (2012).

42. Howe, K. et al. The zebrafish reference genome sequence and its relationship to the human genome. Nature 496, 498–503 (2013).

43. White, R.J. et al. A high-resolution mRNA expression time course of embryonic development in zebrafish. Elife 6(2017).

44. Lawson, N.D. et al. An improved zebrafish transcriptome annotation for sensitive and comprehensive detection of cell type-specific genes. Elife 9(2020).

45. El-Brolosy, M.A. et al. Genetic compensation triggered by mutant mRNA degradation. Nature 568, 193–197 (2019).

46. Paczkowska, M. et al. Integrative pathway enrichment analysis of multivariate omics data. Nat Commun 11, 735 (2020).

47. Buenrostro, J.D., Giresi, P.G., Zaba, L.C., Chang, H.Y. & Greenleaf, W.J. Transposition of native chromatin for fast and sensitive epigenomic profiling of open chromatin, DNA-binding proteins and nucleosome position. Nature Methods 10, 1213–1218 (2013).

48. Ernst, J. & Kellis, M. ChromHMM: automating chromatin-state discovery and characterization.Nature Methods 9, 215–216 (2012).

49. Ernst, J. & Kellis, M. Chromatin-state discovery and genome annotation with ChromHMM. Nature Protocols 12, 2478–2492 (2017).

50. van Steensel, B., Fu, Y., Sinha, M., Peterson, C.L. & Weng, Z. The Insulator Binding Protein CTCF Positions 20 Nucleosomes around Its Binding Sites across the Human Genome. PLoS Genetics 4(2008).

51. Andersson, R. et al. An atlas of active enhancers across human cell types and tissues. Nature 507, 455–461 (2014).

52. Crispatzu, G. et al. The chromatin, topological and regulatory properties of pluripotency-associated poised enhancers are conserved in vivo. Nature Communications 12(2021).

53. Tena, J.J. et al. Comparative epigenomics in distantly related teleost species identifies conserved cis-regulatory nodes active during the vertebrate phylotypic period. Genome Research 24, 1075–1085 (2014).

54. Harmston, N. et al. Topologically associating domains are ancient features that coincide with Metazoan clusters of extreme noncoding conservation. Nature Communications 8(2017).

55. Kaaij, L.J.T., van der Weide, R.H., Ketting, R.F. & de Wit, E. Systemic Loss and Gain of Chromatin Architecture throughout Zebrafish Development. Cell Reports 24, 1–10.e4 (2018).

56. Wike, C.L. et al. Chromatin architecture transitions from zebrafish sperm through early embryogenesis. Genome Research 31, 981–994 (2021).

57. Whyte, Warren A. et al. Master Transcription Factors and Mediator Establish Super-Enhancers at Key Cell Identity Genes. Cell 153, 307–319 (2013).

58. Hnisz, D. et al. Super-Enhancers in the Control of Cell Identity and Disease. Cell 155, 934–947 (2013).

59. Villar, D. et al. Enhancer Evolution across 20 Mammalian Species. Cell 160, 554–566 (2015).

60. Xiao, S. et al. Comparative Epigenomic Annotation of Regulatory DNA. Cell 149, 1381–1392 (2012).

61. Crollius, H.R., Gilardi-Hebenstreit, P., Torbey, P. & Clément, Y. Enhancer–gene maps in the human and zebrafish genomes using evolutionary linkage conservation. Nucleic Acids Research 48, 2357–2371 (2020).

62. Engstrom, P.G., Ho Sui, S.J., Drivenes, O., Becker, T.S. & Lenhard, B. Genomic regulatory blocks underlie extensive microsynteny conservation in insects. Genome Research 17, 1898–1908 (2007).

63. Irie, N. & Kuratani, S. Comparative transcriptome analysis reveals vertebrate phylotypic period during organogenesis. Nature Communications 2(2011).

64. Sawilowsky, S.S. New Effect Size Rules of Thumb. Journal of Modern Applied Statistical Methods 8, 597–599 (2009).

65. Pradeepa, M.M. et al. Histone H3 globular domain acetylation identifies a new class of enhancers. Nature Genetics 48, 681–686 (2016).

66. Farrell, J.A. et al. Single-cell reconstruction of developmental trajectories during zebrafish embryogenesis. Science 360(2018).

67. Briggs, J.A. et al. The dynamics of gene expression in vertebrate embryogenesis at single-cell resolution. Science 360(2018).

68. Farnsworth, D.R., Saunders, L.M. & Miller, A.C. A single-cell transcriptome atlas for zebrafish development. Dev Biol 459, 100–108 (2020).

69. Fornes, O. et al. JASPAR 2020: update of the open-access database of transcription factor binding profiles. Nucleic Acids Research (2019).

70. Balwierz, P.J. et al. ISMARA: automated modeling of genomic signals as a democracy of regulatory motifs. Genome Res 24, 869–84 (2014).

71. Davidson, E.H. Emerging properties of animal gene regulatory networks. Nature 468, 911–920 (2010).

72. Housden, B.E. et al. Loss-of-function genetic tools for animal models: cross-species and cross-platform differences. Nature Reviews Genetics 18, 24–40 (2016).

73. Sakaue-Sawano, A. et al. Visualizing Spatiotemporal Dynamics of Multicellular Cell-Cycle Progression. Cell 132, 487–498 (2008).

74. Ma, Z. et al. PTC-bearing mRNA elicits a genetic compensation response via Upf3a and COMPASS components. Nature 568, 259–263 (2019).

75. Celniker, S.E. et al. Unlocking the secrets of the genome. Nature 459, 927–30 (2009).

76. Kodama, Y., Shumway, M., Leinonen, R. & International Nucleotide Sequence Database, C. The Sequence Read Archive: explosive growth of sequencing data. Nucleic Acids Res 40, D54–6 (2012).

77. Ruzicka, L. et al. The Zebrafish Information Network: new support for non-coding genes, richer Gene Ontology annotations and the Alliance of Genome Resources. Nucleic Acids Res 47, D867–D873 (2019).

78. Kimmel, C.B., Ballard, W.W., Kimmel, S.R., Ullmann, B. & Schilling, T.F. Stages of embryonic development of the zebrafish. Developmental Dynamics 203, 253–310 (1995).

79. Murata, M. et al. Detecting Expressed Genes Using CAGE. in Transcription Factor Regulatory Networks 67–85 (2014).

80. Takahashi, H., Lassmann, T., Murata, M. & Carninci, P. 5′ end–centered expression profiling using cap-analysis gene expression and next-generation sequencing. Nature Protocols 7, 542–561 (2012).

81. D’Orazio, F.M. et al. Germ cell differentiation requires Tdrd7-dependent chromatin and transcriptome reprogramming marked by germ plasm relocalization. Developmental Cell 56, 641–656.e5 (2021).

82. Díaz, N. et al. Chromatin conformation analysis of primary patient tissue using a low input Hi-C method. Nature Communications 9(2018).

83. Acemel, R.D. et al. A single three-dimensional chromatin compartment in amphioxus indicates a stepwise evolution of vertebrate Hox bimodal regulation. Nature Genetics 48, 336–341 (2016).

84. Amemiya, H.M., Kundaje, A. & Boyle, A.P. The ENCODE Blacklist: Identification of Problematic Regions of the Genome. Scientific Reports 9(2019).

85. Haberle, V., Forrest, A.R.R., Hayashizaki, Y., Carninci, P. & Lenhard, B. CAGEr: precise TSS data retrieval and high-resolution promoterome mining for integrative analyses. Nucleic Acids Research 43, e51–e51 (2015).

86. Balwierz, P.J. et al. Methods for analyzing deep sequencing expression data: constructing the human and mouse promoterome with deepCAGE data. Genome Biology 10(2009).

87. Nepal, C. et al. Dynamic regulation of the transcription initiation landscape at single nucleotide resolution during vertebrate embryogenesis. Genome Research 23, 1938–1950 (2013).

88. Etard, C. et al. Loss of function of myosin chaperones triggers Hsf1-mediated transcriptional response in skeletal muscle cells. Genome Biology 16(2015).

89. Meier, M. et al. Cohesin facilitates zygotic genome activation in zebrafish. Development (2017).

90. Marlétaz, F. et al. Amphioxus functional genomics and the origins of vertebrate gene regulation.Nature 564, 64–70 (2018).

91. Dobin, A. et al. STAR: ultrafast universal RNA-seq aligner. Bioinformatics 29, 15–21 (2013).

92. Pertea, M., Kim, D., Pertea, G.M., Leek, J.T. & Salzberg, S.L. Transcript-level expression analysis of RNA-seq experiments with HISAT, StringTie and Ballgown. Nature Protocols 11, 1650–1667 (2016).

93. Niknafs, Y.S., Pandian, B., Iyer, H.K., Chinnaiyan, A.M. & Iyer, M.K. TACO produces robust multisample transcriptome assemblies from RNA-seq. Nature Methods 14, 68–70 (2016).

94. Patro, R., Duggal, G., Love, M.I., Irizarry, R.A. & Kingsford, C. Salmon provides fast and bias-aware quantification of transcript expression. Nature Methods 14, 417–419 (2017).

95. Zerbino, D.R. et al. Ensembl 2018. Nucleic Acids Research 46, D754–D761 (2018).

96. Hahne, F. & Ivanek, R. Visualizing Genomic Data Using Gviz and Bioconductor. in Statistical Genomics 335–351 (2016).

97. Irimia, M. et al. Extensive conservation of ancient microsynteny across metazoans due to cis-regulatory constraints. Genome Research 22, 2356–2367 (2012).

98. de la Calle Mustienes, E., Gómez-Skarmeta, J.L. & Bogdanović, O. Genome-wide epigenetic cross-talk between DNA methylation and H3K27me3 in zebrafish embryos. Genomics Data 6, 7–9 (2015).

99. Ulitsky, I., Shkumatava, A., Jan, Calvin H., Sive, H. & Bartel, David P. Conserved Function of lincRNAs in Vertebrate Embryonic Development despite Rapid Sequence Evolution. Cell 147, 1537–1550 (2011).

100. Li, Q., Brown, J.B., Huang, H. & Bickel, P.J. Measuring reproducibility of high-throughput experiments. The Annals of Applied Statistics 5(2011).

101. Bolger, A.M., Lohse, M. & Usadel, B. Trimmomatic: a flexible trimmer for Illumina sequence data. Bioinformatics 30, 2114–2120 (2014).

102. Chen, H., Smith, A.D. & Chen, T. WALT: fast and accurate read mapping for bisulfite sequencing. Bioinformatics 32, 3507–3509 (2016).

103. Tarasov, A., Vilella, A.J., Cuppen, E., Nijman, I.J. & Prins, P. Sambamba: fast processing of NGS alignment formats. Bioinformatics 31, 2032–2034 (2015).

104. Li, H. et al. The Sequence Alignment/Map format and SAMtools. Bioinformatics 25, 2078–2079 (2009).

105. Kent, W.J., Zweig, A.S., Barber, G., Hinrichs, A.S. & Karolchik, D. BigWig and BigBed: enabling browsing of large distributed datasets. Bioinformatics 26, 2204–2207 (2010).

106. Ramírez, F., Dündar, F., Diehl, S., Grüning, B.A. & Manke, T. deepTools: a flexible platform for exploring deep-sequencing data. Nucleic Acids Research 42, W187–W191 (2014).

107. Quinlan, A.R. & Hall, I.M. BEDTools: a flexible suite of utilities for comparing genomic features. Bioinformatics 26, 841–842 (2010).

108. Schep, A.N. et al. Structured nucleosome fingerprints enable high-resolution mapping of chromatin architecture within regulatory regions. Genome Research 25, 1757–1770 (2015).

109. McInnes, L., Healy, J., Saul, N. & Großberger, L. UMAP: Uniform Manifold Approximation and Projection. Journal of Open Source Software 3(2018).

110. Thodberg, M., Thieffry, A., Vitting-Seerup, K., Andersson, R. & Sandelin, A. CAGEfightR: analysis of 5’-end data using R/Bioconductor. BMC Bioinformatics 20, 487 (2019).

111. Tan, G. & Lenhard, B. TFBSTools: an R/bioconductor package for transcription factor binding site analysis. Bioinformatics 32, 1555–1556 (2016).

112. Engström, P.G., Fredman, D. & Lenhard, B. Ancora: a web resource for exploring highly conserved noncoding elements and their association with developmental regulatory genes. Genome Biology 9(2008).

113. Hubisz, M.J., Pollard, K.S. & Siepel, A. PHAST and RPHAST: phylogenetic analysis with space/time models. Briefings in Bioinformatics 12, 41–51 (2010).

114. Chen, Z. et al. De novo assembly of the goldfish (Carassius auratus) genome and the evolution of genes after whole-genome duplication. Science Advances 5(2019).

115. Lawrence, M., Gentleman, R. & Carey, V. rtracklayer: an R package for interfacing with genome browsers. Bioinformatics 25, 1841–1842 (2009).

116. Akalin, A., Franke, V., Vlahovi ek, K., Mason, C.E. & Schubeler, D. genomation: a toolkit to summarize, annotate and visualize genomic intervals. Bioinformatics 31, 1127–1129 (2014).

117. Kohonen, T. Self-organized formation of topologically correct feature maps. Biological Cybernetics 43, 59–69 (1982).

118. Wehrens, R. & Kruisselbrink, J. Flexible Self-Organizing Maps in kohonen 3.0. Journal of Statistical Software 87(2018).

119. Zhang, Y. et al. Model-based Analysis of ChIP-Seq (MACS). Genome Biology 9(2008).

120. Kruse, K., Hug, C.B. & Vaquerizas, J.M. FAN-C: a feature-rich framework for the analysis and visualisation of chromosome conformation capture data. Genome Biology 21(2020).

121. Kruse, K., Hug, C.B., Hernández-Rodríguez, B. & Vaquerizas, J.M. TADtool: visual parameter identification for TAD-calling algorithms. Bioinformatics 32, 3190–3192 (2016).

122. Nash, A.J., Lenhard, B. & Birol, I. A novel measure of non-coding genome conservation identifies genomic regulatory blocks within primates. Bioinformatics 35, 2354–2361 (2019).

123. Wingett, S. et al. HiCUP: pipeline for mapping and processing Hi-C data. F1000Res 4, 1310 (2015).

124. Heinz, S. et al. Simple combinations of lineage-determining transcription factors prime cis-regulatory elements required for macrophage and B cell identities. Mol Cell 38, 576–89 (2010).

125. Gu, Z., Eils, R. & Schlesner, M. Complex heatmaps reveal patterns and correlations in multidimensional genomic data. Bioinformatics 32, 2847–2849 (2016).

126. Durinck, S. et al. BioMart and Bioconductor: a powerful link between biological databases and microarray data analysis. Bioinformatics 21, 3439–3440 (2005).

127. Durinck, S., Spellman, P.T., Birney, E. & Huber, W. Mapping identifiers for the integration of genomic datasets with the R/Bioconductor package biomaRt. Nature Protocols 4, 1184–1191 (2009).

128. Dijkstra, E.W. A note on two problems in connexion with graphs. Numerische Mathematik 1, 269–271 (1959).

129. Gorkin, D.U. et al. An atlas of dynamic chromatin landscapes in mouse fetal development.Nature 583, 744–751 (2020).

130. Zhang, T., Zhang, Z., Dong, Q., Xiong, J. & Zhu, B. Histone H3K27 acetylation is dispensable for enhancer activity in mouse embryonic stem cells. Genome Biol 21, 45 (2020).

